# *Mycobacterium tuberculosis* CrgA Forms a Dimeric Structure with Its Transmembrane Domain Sandwiched between Cytoplasmic and Periplasmic β-Sheets, Enabling Multiple Interactions with Other Divisome Proteins

**DOI:** 10.1101/2024.12.05.627054

**Authors:** Yiseul Shin, Ramesh Prasad, Nabanita Das, Joshua A. Taylor, Huajun Qin, Wenhao Hu, Yan-Yan Hu, Riqiang Fu, Rongfu Zhang, Huan-Xiang Zhou, Timothy A. Cross

**Affiliations:** Department of Chemistry and Biochemistry, Florida State University, Tallahassee, FL 32306; National High Magnetic Field Laboratory, Tallahassee, FL 32310; Department of Chemistry, University of Illinois Chicago, Chicago, IL 60607; Institute of Molecular Biophysics, Florida State University, Tallahassee, FL 32306; Department of Physics, University of Illinois Chicago, Chicago, IL 60607

## Abstract

CrgA is a key transmembrane (TM) protein in the cell division process of *Mycobacterium tuberculosis* (*Mtb*), the pathogen responsible for tuberculosis. While many of the *Mtb* divisome proteins have been identified, their structures and interactions remain largely unknown. Previous studies of CrgA using oriented-sample solid-state NMR have defined the tilt and rotation of the TM helices, but the cytoplasmic and periplasmic domains and even the oligomeric state were uncharacterized. Here, combining oriented-sample and magic-angle spinning solid-state NMR spectra, we solved the full-length structure of CrgA. The structure features a dimer with a TM domain sandwiched between a cytoplasmic β-sheet and a periplasmic β-sheet. The β-sheets stabilize dimerization, which in turn increases CrgA’s ability to participate in multiple protein interactions. Within the membrane, CrgA binds FtsQ, CwsA, PbpA, FtsI, and MmPL3 via its TM helices; in the cytoplasm, Lys23 and Lys25 project outward from the β-sheet to interact with acidic residues of FtsQ and FtsZ. The structural determination of CrgA thus provides significant insights into its roles in recruiting other divisome proteins and stabilizing their complexes for *Mtb* cell wall synthesis and polar growth.

## Introduction

CrgA (Rv0011c) is a key transmembrane (TM) protein component of the cell division machinery, or divisome, in *Mycobacterium tuberculosis* (*Mtb*) ^1,2^, the bacterial causative agent for tuberculosis (TB). TB is among the top 10 causes of death, claiming 1.3 million lives per year^3^. Treating drug-susceptible TB requires several months on a combination of antimicrobial drugs ^4,5^. Due to the challenges associated with such complex and lengthy treatment, multidrug-resistant and extensive drug-resistant TB have become increasingly common. A hallmark of TB is its latency, in which *Mtb* remains non-proliferative in the patient’s granulomas ^6,7^. This dormant state can last for decades before activation when the patient’s immune system is impaired due to aging or other diseases. A quarter of the global population is latently infected with *Mtb* ^8^; in the United States, up to 13 million people are living with a latent TB infection ^9^. Latency is the primary reason for the lengthy treatment and drug resistance. It is thus of enormous therapeutic interest to understand the mechanism for the *Mtb* latent state. The interactions of divisome proteins play a key role in establishing this latency ^10–12^. In addition, cell division has emerged as a new antibiotic target to counter multidrug-resistant pathogens ^13,14^. A better understanding of the *Mtb* cell division process through characterizing the structures and interactions of CrgA and other divisome proteins is critical for designing resistance-breaking therapeutics.

The divisome is a dynamic supramolecular complex, with some components constantly present but others appearing at different stages of the cell division process. The scaffold of the divisome is the Z-ring, a ring-like structure formed by the polymerization of FtsZ. In *Mtb*, more than 30 other divisome proteins have been identified, but their spatial arrangements and temporal relations are largely unknown ^1,2,15^. CrgA orchestrates cascades of protein-protein interactions throughout the cell division process. It directly interacts with at least six divisome proteins (Figure 1) ^16–18^. Its binding with FtsZ in the cytoplasm enables its localization to the division site. Within the inner membrane, the binding with FtsQ probably facilitates CrgA’s recruitment of the transpeptidases PbpA and FtsI (also known as PbpB) to the nascent division site for peptidoglycan synthesis. Moreover, the binding of CrgA with MmpL3, the exporter of the main component, mycolic acid, of the outer membrane, ensures coordinated construction of the different layers of the cell wall at the division site. Lastly, the binding of CrgA with CwsA enables the recruitment of Wag31 to the new cell pole for polar growth.

**Figure 1.**
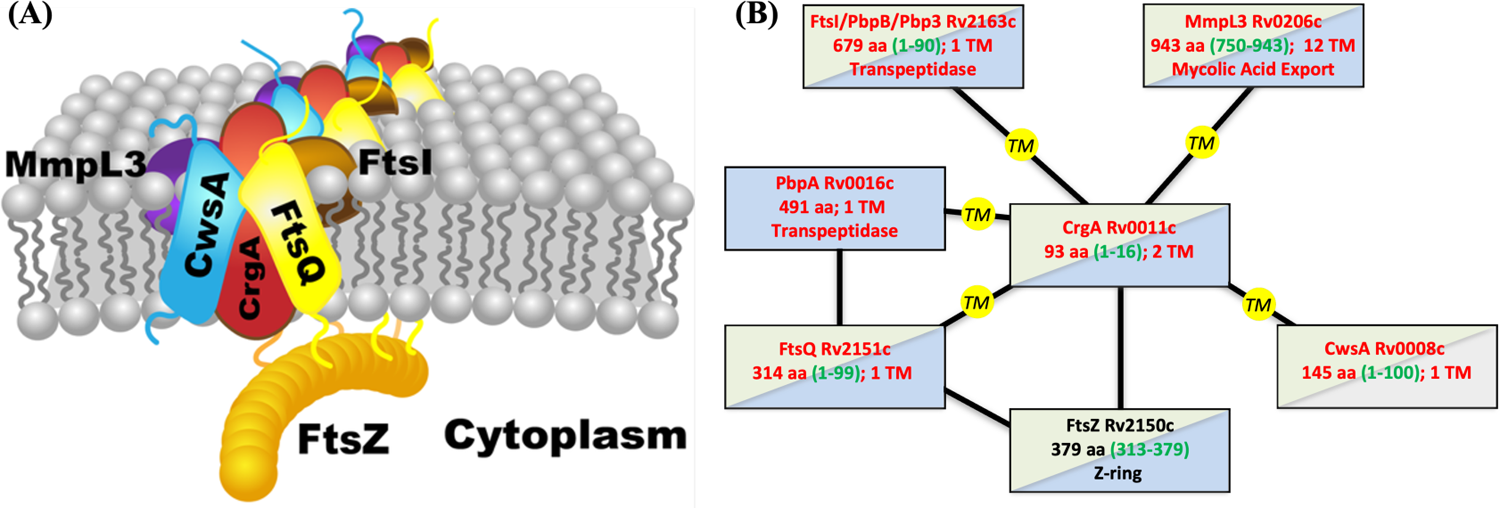
Interactions of CrgA with other *Mtb* divisome proteins. (A) Cartoon representation. (B) Schematic. TM proteins CwsA, FtsQ, FtsI, MmpL3, and PbpA [not depicted in (A)] interact with CrgA through their TM domains, but water-soluble protein FtsZ is suggested to interact with CrgA’s cytoplasmic domain. In (B), TM and water-soluble proteins are indicated by red and black letters, respectively; blue shading indicates structures have been determined for at least a fragment whereas gray shading indicates no such structures; green shading indicates significant disordered regions are present, with residue numbers given by green letters inside parentheses.

*Mtb* CrgA consists of 93 amino acids that form two TM helices (TM1 and TM2; Figure 2A). An initial structural characterization using oriented-sample solid-state NMR spectroscopy (OS ssNMR) precisely defined the tilt angles, 13° for both TM helices, and the rotational orientation of each helix ^19^. The oligomeric state was not known at that time and was assumed to be a monomer. Due to this mistaken assumption and the use of ambiguous distance restraints from magic-angle-spinning (MAS) ssNMR, the packing between the two TM helices in that study is incorrect. In addition to the two TM helices, there is a 28-residue N-terminal region preceding TM1, a 17-residue linker (residues 56-72) between TM1 and TM2, and a single residue following TM2. Both termini are cytoplasmic whereas the interhelical loop is periplasmic; no structural information has been reported for either the cytoplasmic or periplasmic domain. The extreme N-terminal ∼16 residues were found to be disordered whereas the remainder of the N-terminal region and the loop formed either α-helices or β-strands lying parallel to the membrane surface ^19^. A β-hairpin was suggested for the loop.

**Figure 2.**
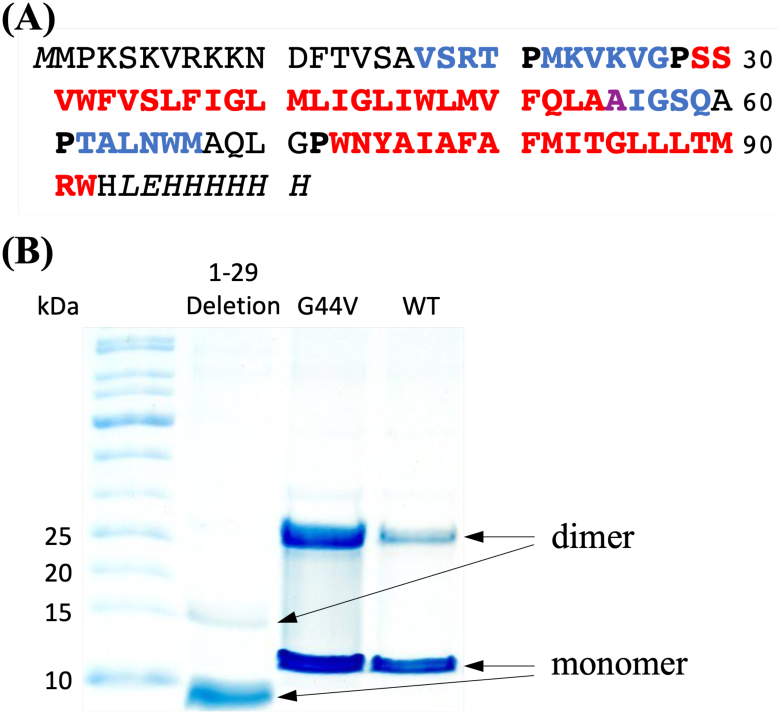
Sequence and oligomeric state of CrgA. (A) Sequence of the expressed CrgA, with TM α-helices in red and β-strands in blue. Ala55 is special in that its NH participates in TM1 hydrogen bonding but its CO participates in β3-β4 hydrogen bonding. Pro residues demarcating secondary structures are in bold. The nonnative His tag starts with LEH. (B) SDS-PAGE gels. Lanes: molecular weight ladder; 1-29 deletion mutant; G44V mutant; and the wild-type (WT) protein.

We recently discovered that CrgA is a dimer and consequently undertook the task of determining the full dimeric structure by ssNMR. By using amino-acid-specific ^13^C labeling along with selective mutations in MAS ssNMR, we obtained interhelical distance restraints both within a monomer and between the monomers in a dimer. We took a similar approach to assign the ^13^C isotropic chemical shifts of residues in the structured portion of the N-terminal region and determined that they form β-strands. Based on this understanding, we reassessed the previously reported OS ssNMR spectra ^19^ to improve the sequence-specific assignment, leading to the conclusion that the structured portion of the N-terminal region and the interhelical loop form β-sheets on the cytoplasmic and periplasmic sides, respectively. Additional distance restraints were obtained to define the relative orientation between the TM helix bundle and the cytoplasmic β-sheet. Finally the dimer structure was refined by running restrained molecular dynamics (MD) simulations in membranes. The structure enables us to examine how CrgA interacts with its various partner proteins and regulates cell division.

## Results

### CrgA forms a dimer with an anticipated β-sheet sandwich structure

The amino-acid sequence of the expressed version of *Mtb* CrgA is presented in Figure 2A, including the nonnative lead residue M0 and C-terminal His tag in italics. The red portion indicates the two TM α-helices, the blue portion represents β-strands, and the black residues are disordered or otherwise outside the regular secondary structures. The lengths of TM1 and TM2 are increased from those defined previously ^19^ by five and one residue, respectively. Importantly, two β-sheets are now identified.

SDS-PAGE gels show that CrgA forms a dimer, and the dimer persists even under the denaturing condition of the gels (Figure 2B). The N-terminal region plays an important role in stabilizing the dimer, as deletion of the first 29 residues significantly weakens the dimer band. We also introduced mutations to probe the dimer interface in the TM domain. G44V, which replaces a single-hydrogen sidechain with a bulkier nonpolar sidechain, increases the intensity of the dimer band, suggesting that Gly44 is located in the intermonomer interface and the bulkier nonpolar sidechain of Val44 enhances helix-helix packing. Likewise, A78V increases the intensity of the dimer band and implicates Ala78 in the intermonomer interface (Figure S1). In contrast, G39V and N74A mutations do not affect the intensity ratio of the dimer and monomer bands, suggesting that these residues face the acyl chains of lipids.

The NMR spectra presented below all report a single set of resonances for each residue, meaning that the CrgA dimer is symmetric. In this symmetric dimer, the termini of the TM α-helices form a parallelogram on both the cytoplasmic and periplasmic sides of the membrane. We anticipate that, on either side of the membrane, these helical termini define the four corners of an intermonomer β-sheet. Below we delineate this unusual dimer structure that has the TM domain sandwiched between two β-sheets.

### Selective intra- and intermonomer restraints produce a range of models for the TM domain

We start by defining the interhelical interfaces in the dimeric TM domain, which consists of two α-helices from each monomer. A widely utilized method for this purpose is dipolar-assisted rotational resonance (DARR) spectroscopy, which detects ^13^C-^13^C correlations between sites with limited dynamics ^20,21^. The strength of these correlations is inversely proportional to the cube of the distance between the nuclei being analyzed. By assessing the correlations through dipolar coupling across a range of mixing times (typically from 50 to 800 ms), the distance between two ^13^C nuclei can be estimated, typically up to a distance of 8 Å. Specifically, at short mixing times (< 100 ms), only intra-residue cross peaks are observable. At longer mixing times (> 300 ms), inter-residue (up to 8 Å) cross peaks can be detected. While revisiting the monomer structure [Protein Data Bank (PDB) entry 2MMU] reported previously ^19^, we found that the interhelical distance restraints were derived from MAS ssNMR spectra of uniformly ^13^C-labeled CrgA with ambiguous assignments. Here we resolved such ambiguity by using amino-acid-specific ^13^C labeling and selective mutations.

We performed MAS ssNMR spectroscopy of CrgA in POPC:POPG liposomes (4:1 molar ratio) to obtain DARR ^13^C-^13^C distance restraints. For interhelical restraints within a monomer, we took advantage of CrgA’s amino-acid sequence to design two distinct ^13^C labeling schemes, one with ^13^C labeling of Tyr and Leu residues and one with ^13^C labeling of Ala and Met residues. The wild-type (WT) CrgA dimer population in the membranes likely coexisted with a sizable monomer population, such that the intensity of any ^13^C-^13^C cross peak would be overwhelmingly from contacts within a monomer instead of between two monomers of a dimer. Our cross peak assignment for the samples with ^13^C labeling of two amino-acid types was thus restricted to residue pairs within a monomer.

For the ^13^C-Tyr/Leu labeling scheme, Tyr was chosen because there is a single such residue, Tyr75, in the entire sequence and this residue falls within TM2. In contrast to Tyr75, Leu is the most abundant amino acid in the TM domain, with six Leu residues in TM1 and three in TM2. The ^13^C-Tyr/Leu labeling scheme not only increases the likelihood of detecting inter-helical interactions by selecting the most prevalent Leu residues across the two helices but also facilitates sequence-specific assignments due to the single Tyr residue Tyr75. This approach resolved the ambiguity problem caused by the high content of hydrophobic residues in CrgA. A Tyr Cα-Leu Cγ cross peak was observed at both 500 ms (Figure 3A) and 800 ms (Figure 3B) mixing times. Within TM2, Tyr75 and the three Leu residues in TM2 are located toward the opposite termini and at least 18 Å apart, so the Tyr Cα-Leu Cγ cross peak must be due to contacts between TM1 and TM2. Since both TM1 and TM2 have very small tilt angles ^19^ and are thus nearly antiparallel to each other, the Leu partner of Tyr75 must be located near the C-terminus of TM1. There are three candidates, Leu45, Leu48, and Leu53, but Leu48 is in the optimal position relative to Tyr75. We therefore assigned this cross peak to a Leu48 Cγ-Tyr75 Cα contact. The cross peak was weak at the 500 ms mixing time and became more prominent at 800 ms (Figure 3C), suggesting that this interhelical contact is at a relatively long distance, which we took as ∼8 Å. For two strictly antiparallel helices, interhelical contacts can only form between two residues that are approximately the same level relative to a membrane surface, and those residue pairs have a constant sum for their residue numbers. In the present case, this sum is 48 + 75 = 123. We thus anticipate that all residue pairs giving rise to interhelical cross peaks to have a sum of residue numbers at or close to 123.

**Figure 3.**
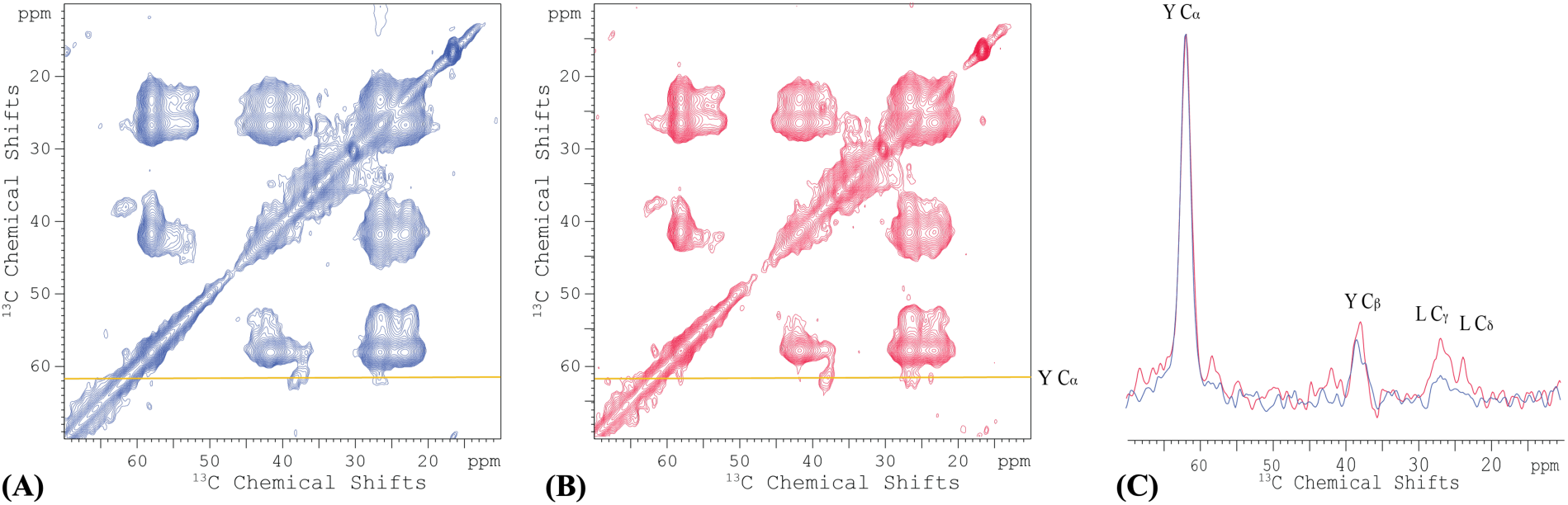
2D DARR ^13^C-^13^C correlation spectra of ^13^C-Tyr/Leu labeled CrgA in POPC:POPG membranes at 265 K. (A) 500 ms mixing time. (B) 800 ms mixing time. A yellow line at 61.7 ppm marks the Tyr Cα frequency. (C) Spectral slices through the Tyr Cα frequency. Spectra were collected at 600 MHz with an 8-kHz spinning rate.

For the ^13^C-Ala/Met labeling scheme, Ala was chosen because it appears at multiple positions in TM2 (Ala76, Ala78, and Ala80). Although Ala54 and Ala55 are on TM1, their terminal position and likely exposure to water may lead to a level of dynamics that precludes strong DARR ^13^C-^13^C cross peaks. Two Met residues, Met41 and Met49, in TM1 as well as Met82 in TM2 could potentially interact with the Ala residues in TM2. A 2D DARR ^13^C-^13^C correlation spectrum with a 500 ms mixing time (Figure S2) revealed two distinct Met Cα-Ala Cα cross peaks near the diagonal. The first cross peak was at (58.3 ppm, 55.5 ppm) and the second was at (60.8 ppm, 55.9 ppm), with a noticeable 2.5 ppm difference between the two Met Cα resonances. These Cα chemical shifts were clearly in the α-helix ranges of Met and Ala ^22^ and thus could be assigned to residues in TM1 and TM2. Met82, situated at *i* - 2 to Ala80 and *i* - 4 to Ala78 in the same TM2 helix, likely accounted for one of the observed Met Cα-Ala Cα cross peaks. The second cross peak could be attributed to an interhelical pair, either between Met41 and Ala80 or between Met49 and Ala76. To assign the Met Cα-Ala Cα cross peaks unambiguously, additional 2D DARR spectra with 100 ms and 800 ms mixing times were recorded (Figure S3A, B). For better comparison, we overlay in Figure S3C the 1D slices through the first and second cross peaks at the different mixing times (100, 500, and 800 ms), with the diagonal peak intensities scaled to the same height. The first cross peak was already visible at the 100 ms mixing time and its intensity saturated when the mixing time was increased to 500 ms, implicating a contact at a relatively short distance; we therefore assigned this cross peak to the intrahelical Met82-Ala78/Ala80 pair. The second cross-peak was not apparent at the 100 ms mixing time, but became discernible at 500 ms and further increased in intensity at 800 ms, indicating a contact at a longer distance (∼8 Å) and thus an interhelical pair.

To assign the second cross peak to either the Met41-Ala80 pair or the Met49-Ala76 pair, a double mutant, A78V/A80G, was designed to eliminate two of the Ala residues in TM2. A DARR ^13^C-^13^C correlation experiment was repeated under the same conditions with the ^13^C-Met/Ala labeled double mutant at a 500 ms mixing time. The 1D slices from 2D DARR spectra of WT CrgA and the double mutant are overlaid in Figure S3D. The elimination of the two Ala residues resulted in the loss of both Met Cα-Ala Cα cross peaks, suggesting that Ala78 and Ala80 accounted for both cross peaks. We thus concluded that the Met49-Ala76 pair could not give rise to a cross peak and the observed second cross peak was from the interhelical Met41 Cα-Ala80 Cα contact. The sum of residue numbers for this pair is 121, close to the number 123 noted above. A range of monomer models, with the TM1-TM2 crossing angle from 7° to 22°, satisfied both these distance restraints from the Leu48 Cγ-Tyr75 Cα and Met41 Cα-Ala80 Cα pairs and the published orientational restraints ^19^. These models will be referred to as acceptable monomer models.

To obtain intermonomer distance restraints within the CrgA dimer, we departed from the design for the intrahelical counterparts in two ways. The first was to label two samples, each with ^13^C labeling on a single type of amino acid, and mix them at a 1:1 molar ratio to form dimers. In this way, ^13^C-^13^C cross peaks between the two types of amino acids could form not within a monomer but only between monomers in a dimer. Note that if an intermonomer residue pair formed a contact, there was only a 25% probability that both partner sites were occupied by ^13^C-labeled residues to generate a cross peak, leading to low sensitivity. To overcome this challenge, we worked with the G44V mutant instead of the WT protein, to increase the dimer stability (Figure 2B) and hence the total dimer population in membranes. In choosing the types of amino acids for ^13^C labeling, we settled on Phe and Met. Phe was chosen because, apart from Phe12, which is disordered and therefore cannot participate in intermonomer interactions or otherwise generate detectable DARR ^13^C-^13^C cross peaks, it is exclusively present in the TM helices, specifically, Phe33, Phe37, and Phe51 in TM1, and Phe79 and Phe81 in TM2. The distribution of Phe residues throughout the TM domain, along with their long aromatic side chains, increases the opportunity for observing intermonomer cross peaks. The second type of amino acid for ^13^C labeling was Met, which is present at eight positions: Met0, Met1, Met22, Met41, Met49, Met67, Met82, and Met90, but only Met41 and Met49 on TM1 and Met82 and Met90 on TM2 can form intermonomer contacts within the TM domain.

The 2D DARR ^13^C-^13^C correlation spectrum of the 1:1 mixture of ^13^C-Phe labeled and ^13^C-Met labeled G44V mutant showed two cross peaks at 500 ms mixing time, located at (131.1 ppm, 33.4 ppm) and (131.1 ppm, 17.4 ppm) (Figure 4). The 131.1 ppm chemical shift was assigned to Phe Cδ and Cε; due to their small chemical shift difference (∼1 ppm), which fell within the linewidth of the cross peaks, no further distinction between Phe Cδ and Cε could be made. The 33.4 ppm chemical shift was assigned to Met Cβ and Cγ, which also have a small chemical shift difference; the 17.4 ppm chemical shift was assigned to Met Cε. Of all the possible pairs of TM Phe and Met residues, only Phe33 and Met90 have a sum of residue numbers that equals 123. The next closest sum is 120, for the Phe79-Met41 pair. We thus assigned the cross peaks in Figure 4 to the intermonomer Phe33-Met90 pair, but could not rule out a contribution from the intermonomer Phe79-Met41 pair. Our initial modeling focused on the Phe33-Met90 pair; 12 atom pairs could contribute to the cross peaks: Phe33 Cδ1-Met90 Cβ, Phe33 Cδ2-Met90 Cβ, Phe33 Cε1-Met90 Cβ, Phe33 Cε2-Met90 Cβ, Phe33 Cδ1-Met90 Cγ, Phe33 Cδ2-Met90 Cγ, Phe33 Cε1-Met90 Cγ, Phe33 Cε2-Met90 Cγ, Phe33 Cδ1-Met90 Cε, Phe33 Cδ2-Met90 Cε, Phe33 Cε1-Met90 Cε, and Phe33 Cε2-Met90 Cε. All the assigned distance restraints from DARR spectra are listed in Table 1.

**Figure 4.**
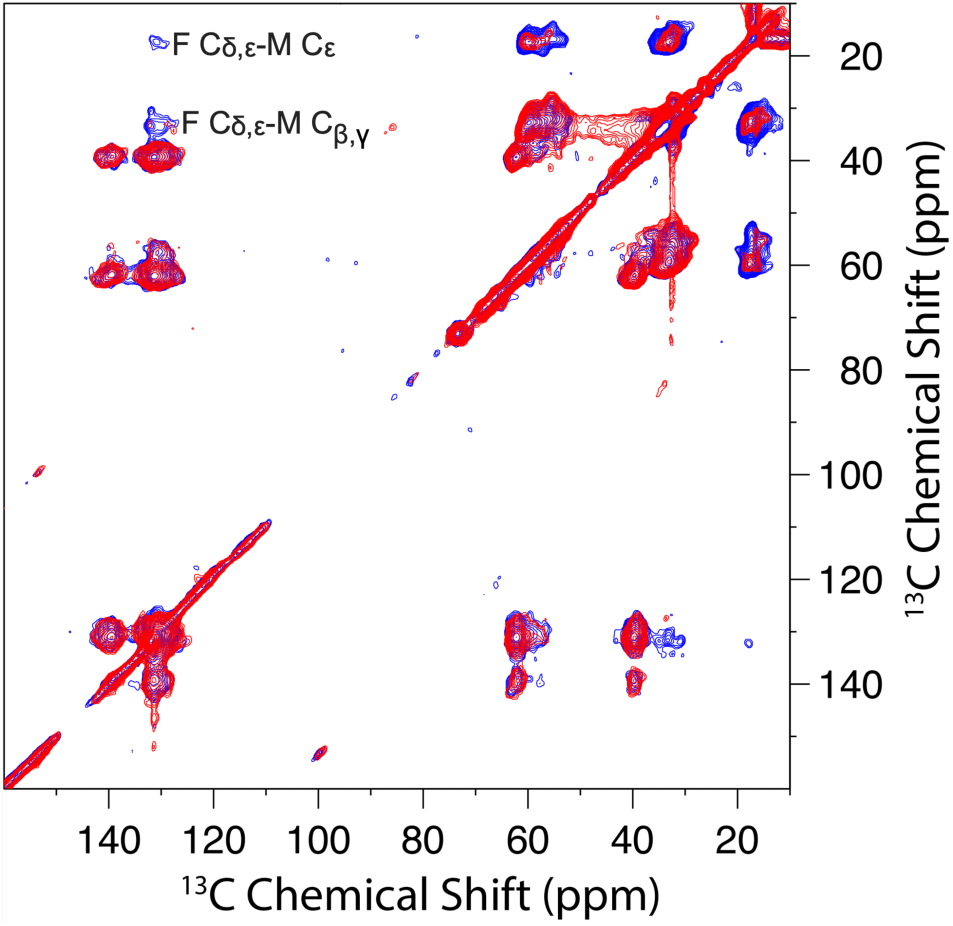
2D DARR ^13^C-^13^C correlation spectra of the 1:1 mixture of ^13^C-Phe labeled and ^13^C-Met labeled CrgA G44V at 50 ms (red) and 500 ms (blue) mixing time and 265 K. Prominent Phe-Met cross peaks were observed at the 500 ms mixing time.

**Table 1.** Intramonomer and intermonomer distance restraints from DARR spectra.

| Intramonomer |  |  |  |
| --- | --- | --- | --- |
| TM1 nucleus | TM2 nucleus | Mixing time (ms) | Distance upper bound (Å) |
| Met41 Cα | Ala80 Cα | 500 & 800 | 8 |
| Leu48 Cγ | Tyr75 Cα | 500 & 800 | 8 |
| Inter monomer |  |  |  |
| Monomer 1 nucleus | Monomer 2 nucleus | Mixing time (ms) | Distance upper bound (Å) |
| Phe33 C$\delta_{1,2}$ | Met90 Cβ | 500 | 8 |
| Phe33 C$\epsilon_{1,2}$ | Met90 Cβ | 500 | 8 |
| Phe33 C$\delta_{1,2}$ | Met90 Cγ | 500 | 8 |
| Phe33 C$\epsilon_{1,2}$ | Met90 Cγ | 500 | 8 |
| Phe33 C$\delta_{1,2}$ | Met90 Cε | 500 | 8 |
| Phe33 C$\epsilon_{1,2}$ | Met90 Cε | 500 | 8 |
| Phe33 C$\delta_{1,2}$ | Val24 Cβ | 600 | 8 |
| Phe33 C$\epsilon_{1,2}$ | Val24 Cβ | 600 | 8 |

We applied C2 symmetry to the acceptable monomer models to generate dimer models. Most of the acceptable monomer models produced dimer models that satisfied the Phe33-Met90 distance restraints. Below we use information from the cytoplasmic and periplasmic domains to narrow the choice of dimer models.

### The N-terminal 16 residues are disordered and exhibit significant membrane association

The first ∼16 residues of CrgA were found to be disordered, based on the lack of significant chemical shift anisotropy in OS ssNMR ^19^. To further characterize the disordered portion, we acquired a 2D^13^C-^13^C TOBSY MAS ssNMR spectrum of uniformly ^13^C labeled CrgA in POPC:POPG liposomes (Figure 5A). In contrast to DARR, TOBSY is sensitive to dynamic residues instead of rigid ones.

**Figure 5.**
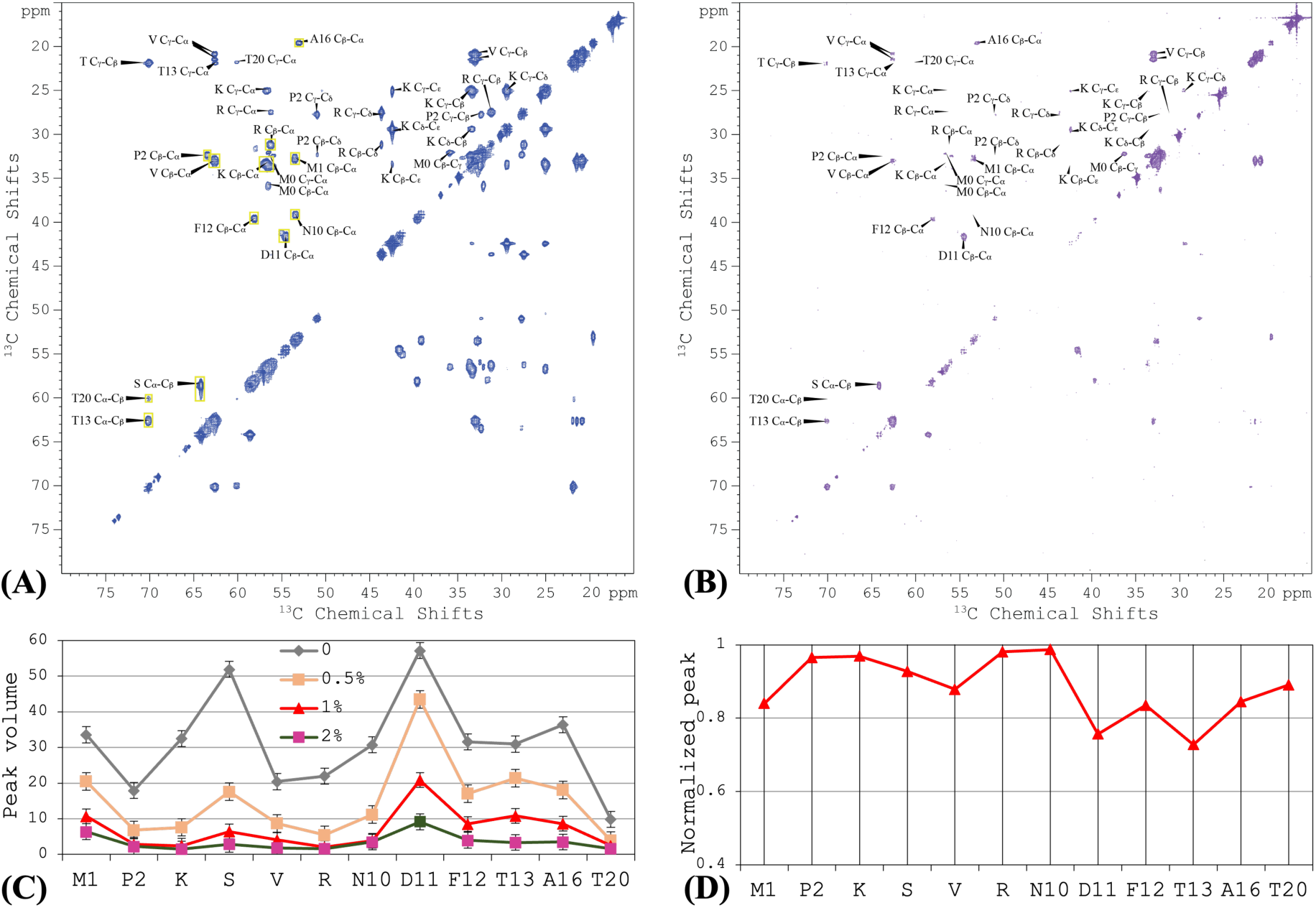
2D TOBSY ^13^C-^13^C correlation spectra of uniformly ^13^C-labeled CrgA in POPC:POPG liposomes without and with 16:0 PE-DTPA (Gd^3+^). (A) Without 16:0 PE-DTPA (Gd^3+^). The yellow rectangles show the areas used to integrate the Cα-Cβ cross peak of each amino acid. (B) With 1% 16:0 PE-DTPA (Gd^3+^). (C) Peak volumes for each residue, from TOBSY spectra in POPC:POPG liposomes doped with 0, 0.5%, 1%, 2% of 16:0 PE-DTPA (Gd^3+^). (D) Normalized peak volume, (1 – *I*1/*I*0)/(1 – *I*2/*I*0), to indicate propensity of membrane association. Spectra were collected at 800 MHz using a 12-kHz spinning rate at 300 K.

We were able to assign a subset of the cross peaks to 9 of the N-terminal 20 residues; other cross peaks were only assigned to amino-acid types, including Lys, Ser, Val, and Arg. The assignment was aided by TOBSY spectra collected on mutants with residue 2-9 deletion or V6/14/17L, T20A, or T20A/M22I substitutions. The TOBSY spectra showed that the N-terminal disordered portion extended to at least Ala16. Indeed, all 11 types of amino acids (Met, Pro, Lys, Ser, Val, Arg, Asn, Asp, Phe, Thr, and Ala) in the N-terminal 16 residues appeared in the TOBSY spectrum of WT CrgA. By contrast, no other amino-acid types appeared in the spectrum, such as Ile, Gly, Gln, Leu, and Trp that are present in the interhelical loop or His and Glu in the C-terminus, suggesting that the disorder is limited to the N-terminal region. However, the next 10 residues beyond Ala16 consist of 7 of the 11 types of amino acids in the first 16 residues, so there was no clear indication of the boundary between disordered and structured portions, except that a weak Cα-Cβ cross peak for Thr20 in the TOBSY spectrum suggested limited dynamics for this site. Moreover, the Thr20 Cα chemical shift, 60.0 ppm, was upfield of the Thr13 counterpart; the latter, at 62.6 ppm, is in the random-coil range ^22^, so the upfield chemical shift implicates Thr20 in a β-strand. In short, the disordered portion of the N-terminal region likely terminates before Thr20.

The inner membrane of *Mtb* is highly enriched in acidic lipids ^23^; the lipid composition, POPC:POPG at 4:1 ratio, was partly chosen to reflect the acidic nature of the inner membrane. On the other hand, 5 of the first 10 residues of CrgA are positively charged, suggesting strong potential for membrane association ^24^. We probed the membrane association of the N-terminal disordered region based on paramagnetic relaxation enhancement (PRE), by doping the membranes with a lipid, 1,2-dipalmitoyl-sn-glycero-3-phosphoethanolamine-N-diethylenetriaminepentaacetic acid, that had a spin label Gd^3+^ chelated to its headgroup [16:0 PE-DTPA (Gd^3+^)]. Residues that came close to the spin label would experience PRE, leading to reduced NMR signals. Indeed, with increasing levels of 16:0 PE-DTPA (Gd^3+^), cross peaks in the TOBSY spectra decreased in intensity (Figure 5B). We quantified the PRE effects by integrating the Cα-Cβ peak intensity for each residue (yellow boxes in Figure 5A); for Cα-Cβ cross peaks assigned only to an amino-acid type, the peak volume was divided by the number of residues in that amino-acid type among the N-terminal 20 residues (Figure 5C). At 1% 16:0 PE-DTPA (Gd^3+^), the cross peaks of Pro2, Lys, Arg, and Asn10 all but disappeared (Figure 5B, C), indicating significant membrane association. We further used the parameter, (1 – *I*_1_/*I*_0_)/(1 – *I*_2_/*I*_0_) where *I_x_* represents the peak volume at *x*% 16:0 PE-DTPA (Gd^3+^), as a measure of the membrane association propensity (Figure 5D), which shows that residues D_11_FT_13_--A_16_ and perhaps also the intervening V_14_S_15_ have lower membrane-association propensities than the first 10 residues.

### The rest of the N-terminal region forms an intermonomer β-sheet

To characterize the structured portion of the N-terminal region, we used ^13^C-^13^C correlation spectra to assign ^13^C isotropic chemical shifts. We first chose Lys for ^13^C labeling because Lys is present only in the N-terminal region, in the disordered portion (Lys3, Lys5, Lys8, and Lys9) and also in the potentially structured portion (Lys23 and Lys25). The ^13^C-^13^C correlation spectrum of ^13^C-Lys labeled CrgA (Figure S4) showed Cα-Cβ, Cα-CO, and Cβ-CO cross peaks with random-coil values, (56.1 ppm, 32.8 ppm, 176.1 ppm), for (Cα, Cβ, CO) chemical shifts as well as Cα-Cβ and Cα-CO cross peaks with (Cα, Cβ, CO) chemical shifts at (54.6 ppm, 35.2 ppm, 174.1 ppm) that are consistent with β-strands, with upfield Cα and CO chemical shifts and a downfield Cβ chemical shift relative to random-coil values. α-Helical residues would have chemical-shift changes in the opposite directions. The second set of cross peaks thus indicated that Lys23 and Lys25 are located in a β-strand, likely delimited by Pro21 and Pro28. Cα-Cγ and Cα-Cδ cross peaks of the β-strand Lys residues were also detected.

Our second ^13^C-^13^C correlation spectrum was from ^13^C-Val labeled CrgA (Figure S5, black). Val appears in both the N-terminal region (Val6, Val14, Val17, Val24, and Val26) and TM1 (Val31, Val34, and Val50). Val6 and Val14 are in the disordered portion whereas Val24 and Val26 are in the structured portion of the N-terminal region; where Val17 fell was uncertain. Correspondingly, the ^13^C-^13^C correlation spectrum showed three distinct Cα-Cβ cross peaks. A strong, broad cross peak, with Cα chemical shift spanning 63-68 ppm and Cβ chemical shift spanning 30-33 ppm, was assigned to the TM1 Val residues. Indeed, in the ^13^C-^13^C correlation spectrum of a ^13^C-Val labeled mutant in which residues 2-28 were deleted, only this Cα-Cβ cross peak remained (Figure S5, red). Of the remaining two Cα-Cβ cross peaks, one, centered at Cα ∼ 62 ppm and Cβ ∼ 33 ppm, was assigned to disordered Val residues in the N-terminal region; the last cross peak, centered at Cα ∼ 60.5 ppm and Cβ ∼ 34.5 ppm, thus belonged to the structured Val residues in the N-terminal region. The latter chemical shifts, upfield for Cα and downfield for Cβ relative to random-coil values, place these Val residues in β-strands. Therefore, Val24, and Val26 and possibly even Val17 are located in a β-sheet. Here again, a prominent Cα-Cγ cross peak of the β-strand Val residues was detected. By now there is strong evidence for K_23_VKV_26_ belonging to a β-strand delimited by Pro21 and Pro28.

To confirm that Met22 is also part of this β-strand, we collected a ^13^C-^13^C correlation spectrum for ^13^C-Met labeled CrgA (Figure S6A, black). To assign the cross peaks of Met22, we also collected the corresponding spectrum for the T20A/M22I mutant. A Cα-Cβ cross peak in the WT spectrum that was missing in the mutant spectrum (Figure S6A, red) could be assigned to Met22. Its Cα chemical shift of ∼ 54 ppm and Cβ chemical shift of ∼ 36.4 ppm deviate from the random-coil values (∼55.4 ppm and 33.7 ppm) in the upfield and downfield directions, respectively, thereby confirming Met22 as a β-strand residue. Moreover, ^13^C-Ile labeling of these two constructs (Figure S6B) showed that the Cα-Cβ cross peak of Ile22 in the mutant also had deviations of Cα and Cβ chemical shifts in the directions expected for a β-strand residue.

Together, the isotropic chemical shifts in ^13^C-^13^C correlation spectra showed that the residues between Pro21 and Pro28 form a β-strand. Our gel results revealed that the N-terminal region is involved in the dimer formation of CrgA (Figure 2B). More specifically, dimer formation in the cytoplasm is most likely mediated by the structured portion of the N-terminal region. Because M_22_KVKVG_27_ form a β-strand, the pairing between two such β-strands, one from each monomer, into a β-sheet is then the simplest mechanism for dimerization. V_17_SRT_20_, between the disordered Ala16 and Pro21, may form an additional β-strand to reinforce the cytoplasmic β-sheet. This point is supported by the upfield Cα chemical shift of Thr20 Cα relative to the random-coil value in the TOBSY spectrum (Figure 5A).

With the foregoing new understanding of the N-terminal region, we reassessed the OS ssNMR spectra reported previously ^19^ to complete the sequence-specific assignments for the nonhelical residue. This reassessment also resulted in modest extensions of the TM helices.

These OS ssNMR spectra correlate the dipolar coupling (DC) of backbone ^15^N-^1^H and the anisotropic chemical shift (aCS) of ^15^N. We collected OS ssNMR spectra on 12 CrgA samples with amino-acid-specific ^15^N labeling (Figure 6 and Figure S7). Except for His and Gln residues, all the structured residues having a protonated amide nitrogen were observed by OS ssNMR. The new assignments of the cross peaks are listed in Table 2. The backbone amides of the first 16 residues displayed essentially 0 dipolar couplings and isotropic chemical shifts. For example, Ala16 was assigned to a cross peak at 0 dipolar coupling and 129 ppm anisotropic chemical shift (Figure 6A); the latter value is very close to the amide ^15^N isotropic chemical shift of an Ala residue in a random-coil conformation. The first structured segment in the N-terminal region is residues V_17_SRT_20_, followed by the secondary-structure breaker Pro21, and then another secondary structure formed by M_22_KVKVG_27_, again terminated by a Pro residue (Pro28) before entering the first TM helix.

**Figure 6.**
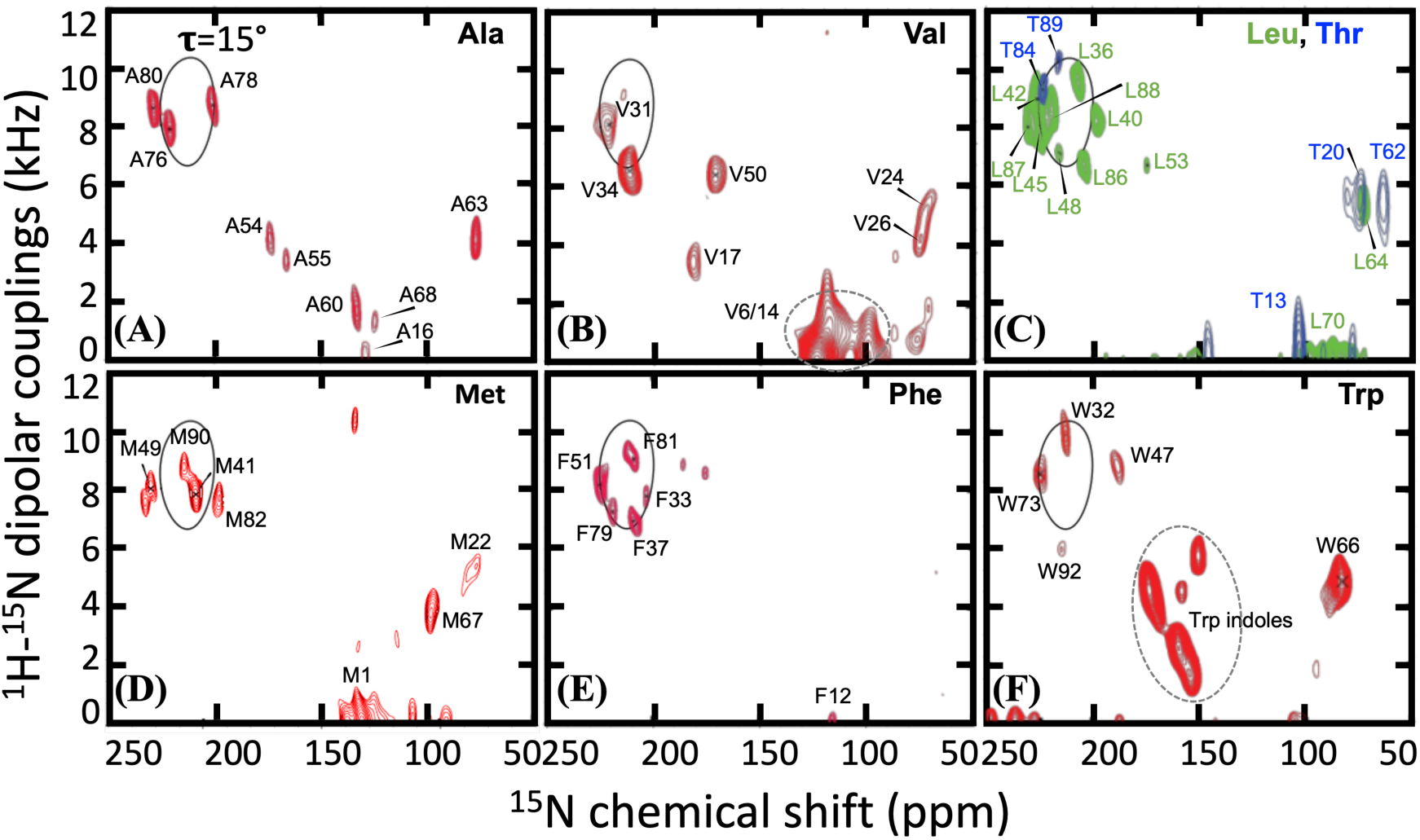
OS ssNMR spectra of amino-acid-specific ^15^N-labeled samples. A PISA wheel for a TM helix with a 15° tilt angle is shown in solid curve. (A) ^15^N-Ala. (B) ^15^N-Val. (C) ^15^N-Leu and ^15^N-Thr. (D) ^15^N-Met. (E) ^15^N-Phe. (F) ^15^N-Trp. Panel (A) and the helical portions of other panels were published previously in ref 19.

**Table 2.** ^15^N-^1^H dipolar coupling (DC in kHz) and ^15^N anisotropic chemical shift (aCS in ppm) for each residue in CrgA determined by OS ssNMR. (A) N-terminal region. (B) TM1. (C) Interhelical loop. (D) TM2. The assignments for many residues in the nonhelical regions were changed from (cyan; 11 total) or added to (yellow; 4 total) those initially reported in ref 19.

| (A) | AA | DC | aCS | (B) | AA | DC | aCS | (C) | AA | DC | aCS | (D) | AA | DC | aCS |
| --- | --- | --- | --- | --- | --- | --- | --- | --- | --- | --- | --- | --- | --- | --- | --- |
|  | Met1 | 0 | 133 |  | Ser29 | 4.0 | 179 |  | Ile56 | 5.8 | 78 |  | Trp73 | 8.6 | 225 |
|  | Pro2 |  |  |  | Ser30 | 8.5 | 225 |  | Gly57 | 5.1 | 74 |  | Asn74 | 9.6 | 209 |
|  | Lys3 |  |  |  | Val31 | 8.1 | 221 |  | Ser58 | 5-6 | 66-87 |  | Tyr75 | 4.3 | 190 |
|  | Ser4 |  |  |  | Trp32 | 9.8 | 213 |  | Gln59 |  |  |  | Ala76 | 7.9 | 221 |
|  | Lys5 |  |  |  | Phe33 | 7.8 | 204 |  | Ala60 | 1.8 | 134 |  | Ile77 | 10.1 | 220 |
|  | Val6 | 0 | 124 |  | Val34 | 6.6 | 211 |  | Pro61 |  |  |  | Ala78 | 8.7 | 201 |
|  | Arg7 |  |  |  | Ser35 | 9.1 | 228 |  | Thr62 | 5.5 | 62 |  | Phe79 | 7.2 | 219 |
|  | Lys8 |  |  |  | Leu36 | 9.6 | 207 |  | Ala63 | 4.1 | 78 |  | Ala80 | 8.6 | 229 |
|  | Lys9 |  |  |  | Phe37 | 7.0 | 209 |  | Leu64 | 5.5 | 74 |  | Phe81 | 9.0 | 209 |
|  | Asn10 |  |  |  | Ile38 | 7.2 | 222 |  | Asn65 | 5-6 | 66-87 |  | Met82 | 7.8 | 194 |
|  | Asp11 |  |  |  | Gly39 | 9.7 | 220 |  | Trp66 | 5.0 | 85 |  | Ile83 | 6.1 | 219 |
|  | Phe12 | 0 | 116 |  | Leu40 | 8.2 | 198 |  | Met67 | 3.9 | 105 |  | Thr84 | 9.3 | 227 |
|  | Thr13 | 0 | 106 |  | Met41 | 7.9 | 203 |  | Ala68 | 1.1 | 125 |  | Gly85 | 9.9 | 201 |
|  | Val14 | 0 | 124 |  | Leu42 | 8.9 | 226 |  | Gln69 |  |  |  | Leu86 | 6.6 | 204 |
|  | Ser15 |  |  |  | Ile43 | 9.9 | 209 |  | Leu70 | 0 | 97 |  | Leu87 | 7.9 | 230 |
|  | Ala16 | 0 | 129 |  | Gly44 | 9.5 | 193 |  | Gly71 | 1.2 | 120 |  | Leu88 | 8.6 | 219 |
|  | Val17 | 3.5 | 175 |  | Leu45 | 7.8 | 225 |  | Pro72 |  |  |  | Thr89 | 10.2 | 216 |
|  | Ser18 | 5-6 | 66-87 |  | Ile46 | 9.4 | 227 |  |  |  |  |  | Met90 | 8.8 | 207 |
|  | Arg19 | 5.6 | 73 |  | Trp47 | 8.8 | 188 |  |  |  |  |  | Arg91 | 4.7 | 177 |
|  | Thr20 | 5.6 | 74 |  | Leu48 | 7.1 | 216 |  |  |  |  |  | Trp92 | 6.0 | 213 |
|  | Pro21 |  |  |  | Met49 | 8.1 | 222 |  |  |  |  |  |  |  |  |
|  | Met22 | 5.4 | 89 |  | Val50 | 6.4 | 171 |  |  |  |  |  |  |  |  |
|  | Lys23 | 5-6 | 66-87 |  | Phe51 | 8.2 | 225 |  |  |  |  |  |  |  |  |
|  | Val24 | 5.4 | 72 |  | Gln52 |  |  |  |  |  |  |  |  |  |  |
|  | Lys25 | 5-6 | 66-87 |  | Leu53 | 6.6 | 175 |  |  |  |  |  |  |  |  |
|  | Val26 | 4.2 | 74 |  | Ala54 | 4.2 | 173 |  |  |  |  |  |  |  |  |
|  | Gly27 | 5.1 | 78 |  | Ala55 | 3.6 | 166 |  |  |  |  |  |  |  |  |
|  | Pro28 |  |  |  |  |  |  |  |  |  |  |  |  |  |  |

Val17 was assigned to a unique resonance (DC = 3.5 kHz, aCS = 175 ppm) in an ^15^N-Val labeled sample (Figure 6B). The aCS value is well removed from the isotropic chemical shift (∼120 ppm) and separated from the relatively uniform aCS values (∼75 ppm) for Ser18, Arg19, and Thr20. Ser18 along with Lys23 and Lys25 was assigned to a resonance band (5-6 kHz and 66-87 ppm) in the OS ssNMR spectrum of CrgA with reverse labeled ^15^N-Ser/Lys/Asn (Figure S7A), but Arg19 (5.6 kHz, 73 ppm) and Thr20 (5.6 kHz, 74 ppm) could be unambiguously assigned (Figure S7B and Figure 6C). The assignments for the rest of the structured portion of the N-terminal region were: Met22 (5.4 kHz, 89 ppm), Val24 (5.4 kHz, 72 ppm), Val26 (4.2 kHz, 74 ppm), and Gly27 (5.1 kHz, 78 ppm) (Figure 6B, D and Figure S7C). The similar resonance frequencies for Ser18-Thr20 and Met22-Gly27 (DC ∼ 5.5 kHz, aCS ∼ 75 ppm) are in a spectral region that can be attributed to a β-sheet or an α-helix oriented parallel to the bilayer surface and thus perpendicular to the magnetic field in the NMR spectrometer. Only a β-sheet is consistent with the ^13^C-^13^C correlation and TOBSY spectra presented above. This cytoplasmic β-sheet is formed between the two monomers of a symmetric dimer and consists of two β-strands from each monomer, to be named β1 and β2. DARR distance restraints between Phe33 on TM1 and Val24 on the cytoplasmic β-sheet, to be presented below, verified the expected result that the longer β2 strand, formed by M_22_KVKVG_27_, is in the core of the β-sheet and the shorter β1 strand, formed by V_17_SRT_20_ with a frayed Val17, is at the periphery. Note that β1 must extend to Val17, as Ser18’s NH and CO bonds project away from β2 and cannot form interstrand hydrogen bonds but its OS spectral frequencies implicate a β-strand conformation; it falls on Val17 to form interstrand hydrogen bonds with Val24 to keep Ser18 in the β-strand conformation.

The symmetry of the dimer dictates that the two β2 strands of the monomers must align in an antiparallel manner. The two middle residues, Val24 and Lys25, most likely form the fulcrum, which would leave Gly27 and Met22, respectively, as an overhanging residue. We selected the former case, where the Gly27 NH of one monomer can still form a hydrogen bond with the Pro21 CO of the other monomer, thereby explaining the β-strand OS frequencies (5.1 kHz, 78 ppm) of Gly27, but the backbone can bend at this residue to facilitate the connection between β2 and TM1 (Figure S8A). For simplicity, we refer to the two faces of the cytoplasmic β-sheet as upper and lower, with the former toward the membrane hydrophobic core and the latter toward the aqueous phase in the cytoplasm. The nonpolar sidechains of Met22, Val24, and Val26 project from the upper face into the TM helix bundle, whereas the charged sidechains of Lys23 and Lys25 project from the lower face. Within each monomer, the two β-strands are expected to form a β-hairpin. Both Thr20 and Met22 were found to be in β-strand conformations according to ^13^C isotropic chemical shifts, leaving Pro21 as the only non-β-strand residue at the turn between the two β-strands. The resulting turn conforms to type VI ^25,26^, which is observed only infrequently and characterized by a β-strand residue at the *i* + 1 position and a Pro residue at the *i* + 2 position. In the present case, these two positions are occupied by Thr20 and Pro21, respectively; the interstrand hydrogen bonding starts with the Met22-Arg19 pair and ends with the Val24-Val17 pair. This type-VI connected β-hairpin arrangement is corroborated by the fact that the OS ssNMR cross peaks of all residues 18-27 except Pro21 collapse to a small region expected of a β-sheet. Within the β1 strand, Val17 projects from the upper face whereas Ser18 projects from the lower face; Arg19, nominally on the same face of β1 as Val17, can snorkel into the lipid headgroup region.

### A ^13^C-^13^C cross peak fixes the relative orientation and elevation between the TM domain and the cytoplasmic β-sheet

The β2 strand is terminated by Pro28, which is right next to the N-terminal residue Ser29 of TM1. The OS spectral frequencies, (4.0 kHz, 179 ppm), of this Ser residue are very different from the β-sheet values, but not so different from typical TM α-helix values, such as those of Ser30: (8.5 kHz, 225 ppm). Consequently, β2 and TM1 are linked by only two residues: half of Gly27, Pro28, and half of Ser29 (Figure S8A). This short linker places the N-terminus of TM1 on the outer edge of β2, next to the N-terminus of β1. In this position, TM1 blocks β1 from extending beyond Val17 toward the N-terminus of CrgA. The fact that the TM1 N-terminal segments need to be connected with the β2 strands via a short two-residue linker on one side and be next to the N-termini of the β1 strands tightly restricts the rotation of the cytoplasmic β-sheet relative to the TM helix bundle. In addition, Trp32 near the N-terminus of TM1 and Trp92 at the C-terminus of TM2, conveniently serve as markers for ascertaining the relative elevation of the cytoplasmic β-sheet. The benzene rings in the indoles of these Trp residues should have similar elevations as the Val24 and Val26 sidechains, and hence on the upper side of the β-sheet (Figure S8B). Recall that β-sheets are pleated and thus their two faces span a finite thickness. Because

TM1 and TM2 are oriented in opposite directions, the Cα-Cβ bonds of Trp32 and Trp92 also point in opposite directions, with the former toward the β-sheet whereas the latter away from the β-sheet. Thus the Cα atom of Trp92 nearly lies in the plane of the upper face of the β-sheet while the Cα atom of Trp32 is more elevated (toward the membrane center). As Trp92 is its very last residue, the entire TM2 is on the upper side of the cytoplasmic β-sheet. On the other hand, from Trp32, TM1 further extends to Ser29; the higher elevation of Trp32 Cα places Val31 Cα at a similar level as Trp92 Cα and the upper face of the β-sheet. The latter two residue numbers add up to 123, which is the number stated above for TM1 and TM2 residues at the same elevation. TM1 then continues with Ser30 at the same level as the lower face of the β-sheet and terminates with Ser29 crossing into the aqueous phase.

To confirm the orientation and elevation of the cytoplasmic β-sheet relative to the TM domain, ¹³C-Phe and ³C-Val labeled G44V samples were mixed at a 1:1 molar ratio as described above. Val was chosen for labeling due to the absence of Val residues in the interhelical loop, simplifying cross peak assignment. As shown in Figure 7, a single cross peak, indicated by an arrow at 131.2 ppm and 33.2 ppm, was observed at a 600 ms mixing time but not at a 300 ms mixing time, suggesting a distance that is near the upper bound of DARR detection limit (∼8 Å). This cross peak can be assigned to F Cδ,ε-V Cβ. The G44V sequence contains six Phe residues (Phe12, Phe33, Phe37, Phe51, Phe79, and Phe81) and nine Val residues [Val6, Val14, Val17, Val24, Val26, Val31, Val34, Val44 (mutation site), and Val50]. The disordered residues, Phe12, Val6, and Val14, can be eliminated as they would be too dynamic to generate a DARR cross peak. Moreover, the chemical shift of 33.2 ppm is downfield of the random-coil value (∼32.3 ppm) for Val Cβ, and thus belongs to a β-strand residue, which could be Val17, Val24, and Val26. Of the remaining Phe residues, only Phe33 from TM1 is in close proximity to the cytoplasmic β-sheet and thus contributes to this cross peak. Phe33 has already been determined to form intermonomer contacts (Figure 4) and is thus located in the middle of the TM helix bundle. Both Val17 and Val26 are near the edges of the cytoplasmic β-sheet and thus far from the TM1 helix of the other monomer. Val24 is at the center of this β-sheet and thus generates an intermonomer cross peak with Phe33. Therefore, the cross peak in Figure 7 was unambiguously assigned to the intermonomer F33Cδ,ε-V24Cβ pairs.

**Figure 7:**
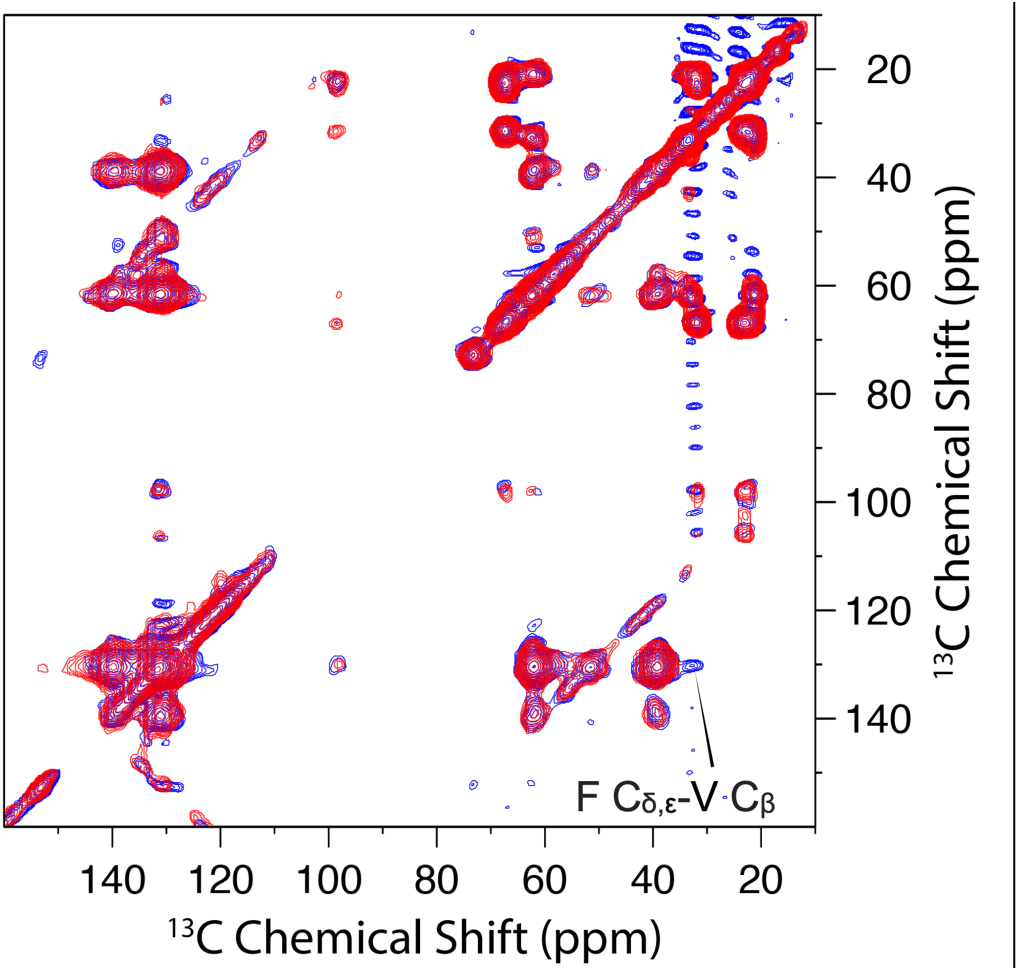
Overlay of 2D DARR ^13^C-^13^C correlation spectra of ^13^C-Phe and ^3^C-Val labeled G44V mutant 1:1 mix at 300 ms (red) and 600 ms (blue) mixing times and 265 K. The cross peak at the 600 ms mixing time indicated by an arrow was attributed to F Cδ,ε-V Cβ and not observed at 300 ms mixing time.

The above-defined positioning of the TM1 N-termini relative to the cytoplasmic β-sheet tightly restricts the intermonomer Ser29-Ser29 Cα-Cα distance to ∼18 Å. Only two of the acceptable dimer models for the TM domain are consistent with this restriction. The two surviving models differ in the distances between the TM1 C-termini. The intermonomer Ala55-Ala55 Cα-Cα distance is ∼22 Å in one model and slightly < 18 Å in the other. The first model will be eliminated below after considering the periplasmic domain, leaving only a single model.

### The interhelical loop also forms an intermonomer β-sheet

We previously suggested that the interhelical loop on the periplasmic side may form a β-hairpin^19^. With the revised assignment of the OS spectra (Table 2) and the new finding that CrgA forms a dimer, we are now certain of β-hairpin formation in each monomer, resulting in a periplasmic β-sheet. The OS assignments allowed us to identify two stretches of β-strand residues, I_56_GSQ_59_ and T_62_ALNWM_67_, to be named β3 and β4, respectively (Figure S9). Interestingly, there are several similarities between the periplasmic and cytoplasmic β-sheets. Here again, the β-hairpin in each monomer involves a turn comprising two residues with a Pro residue, Pro61, at the *i* + 2 position and hence potentially conforming to a type-VI turn; accordingly, hydrogen bonding in the β-hairpin starts with the Thr62 NH-Gln59 CO pair and ends with the Trp66 NH-Ala55 CO pair. This last hydrogen bond is required as without it Ile56 would be frayed from β3, contradicting its β-strand OS frequencies (5.8 kHz, 78 ppm). The Trp66 NH-Ala55 CO hydrogen bond makes Ala55 very special as it is also nominally a part of TM1, with its NH participating in hydrogen bonding there. Also similar to the cytoplasmic β-sheet, the second strand here, β4, is longer than the first strand, β3. We again assigned the longer β4 as the inner strand of the periplasmic β-sheet, for two reasons. First, because TM1 abruptly transitions into β3, with Ala55 engaged in hydrogen bonding in both secondary structural elements, so β3 has to be at the periphery of the parallelogram defined by the periplasmic termini of TM1 and TM2. Second, at the inner position, the longer β4 can form more intermonomer hydrogen bonds than the shorter β3 would.

Similar to the consideration for the cytoplasmic β-sheet, the two middle residues, Leu64 and Asn65, most likely form the fulcrum for the antiparallel alignment of the β4 strands of two monomers, which would leave Met67 and Thr62, respectively, as an overhanging residue. Thr62 is already paired with Gln59 in intramonomer hydrogen bonding, but Met67 has to rely on intermonomer hydrogen bonding to maintain its β-strand conformation; therefore we selected the β4-β4 alignment with Asn65 as the fulcrum (Figure S9). The periplasmic β-sheet tightly restricts the positioning of the TM1 C-termini, because a single residue, Ala55, links them to the β3 strands. The β-sheet with Asn65 at the fulcrum results in a 17 Å intermonomer Ala55-Ala55 Cα-Cα distance, narrowing the choices for the dimer model to a single one.

One last issue to resolve is which face of the periplasmic β-sheet directs toward the membrane interior. Unlike the cytoplasmic β-sheet with nonpolar sidechains projecting from one face and charged sidechains projecting from the opposite face, the periplasmic β-sheet has a mix of polar and nonpolar sidechains projecting from either face, with Ala55, Gly57, Gln59, Thr62, Leu64, and Trp66 on one face and Ile56, Ser58, Ala63, Asn65, and Met67 on the other.

Although the amino-acid composition does not have an apparent bias, there are two good reasons to have the Ala55 face on the membrane side. If the Ile56 face were to be on the membrane side, each Ile56 residue would clash with the preceding TM1 C-terminus. Moreover, each β4-TM2 linker (residues Ala68-Pro72) would have to span the length of the β4 strand to reach TM2, generating potential for clashing with the β-sheet. With the Ala55 face on the membrane side, the TM2 N-terminus is a short distance away from the β4 C-terminus and under the β-hairpin of the other monomer, so the β4-TM2 linker can be easily accommodated at the periphery of the periplasmic β-sheet (Figure S9).

### CrgA forms an unusual dimer structure with the TM domain sandwiched between two β-sheets

We built a dimer model for residues Val17-His93 as described above and placed it in a POPC:POPG membrane to refine it by running restrained MD simulations. The resulting dimer structure, shown in Figure 8, satisfies well both the OS restraints (Figure S10) and DARR restraints. For example, Phe33 of monomer A and Met90 of monomer B have Cε1-Cγ and Cε2-Cε distances ∼ 4.5 Å. Additionally, the sidechains of Met41 and Phe79 have intermonomer distances within 7 Å. That two pairs of residues are within the detectable distance range explains the relatively strong Phe-Met cross peaks in Figure 4.

**Figure 8:**
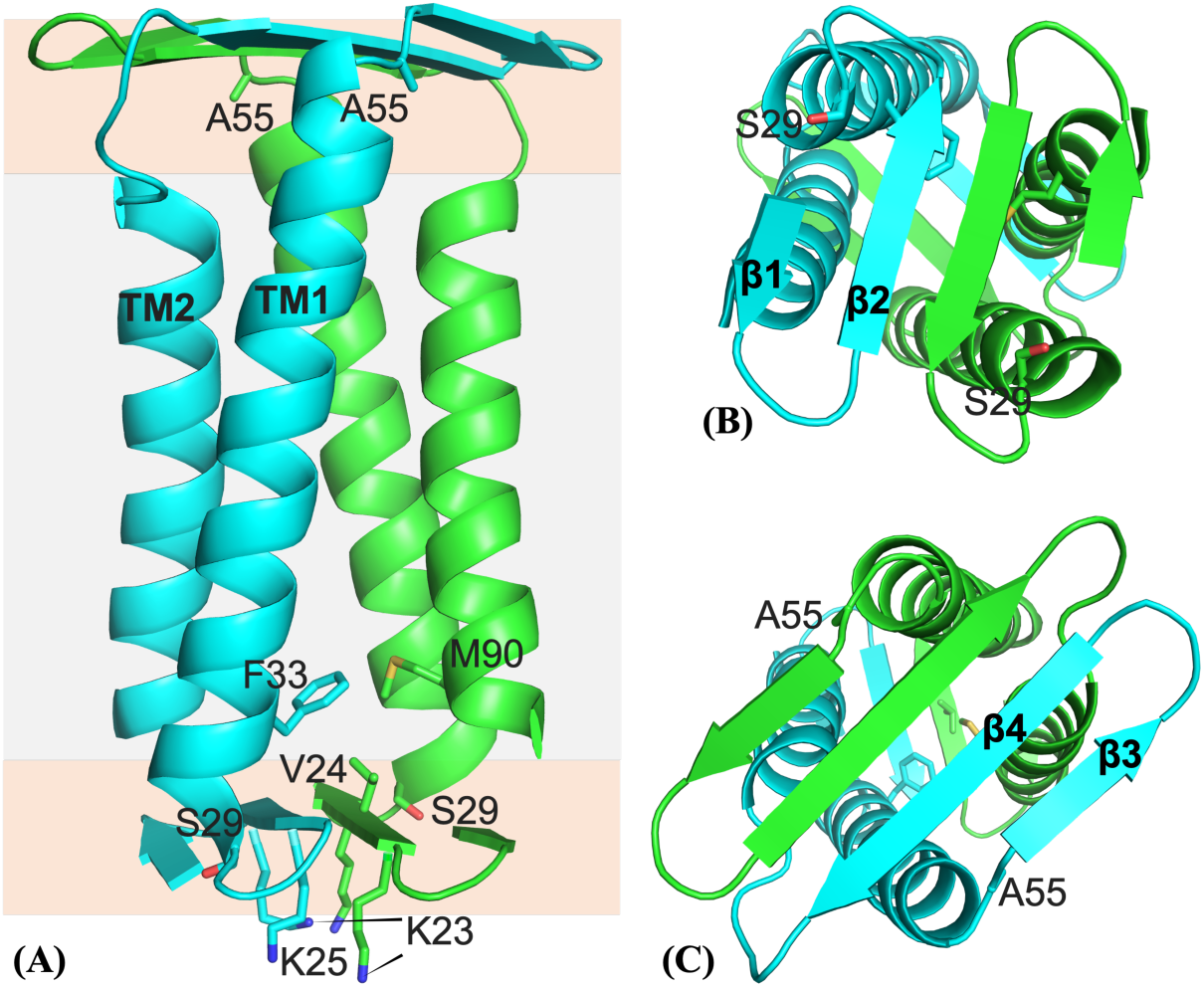
Dimer structure of CrgA. (A) Side view. Gray and orange shading indicate the hydrophobic core and headgroup regions, respectively, of the membrane. (B) Bottom view. (C) Top view.

The CrgA dimer has the TM domain sandwiched between the cytoplasmic β-sheet and the periplasmic β-sheet (Figure 8A). On the cytoplasmic side, the β-hairpin of each monomer is largely free of contact with the TM helices of the other monomer (Figure 8B). By contrast, the periplasmic β-sheet is rotated by 58° relative to the cytoplasmic β-sheet, such that the periplasmic β-hairpin of each monomer is on top of TM2 of the other monomer (Figure 8C). The resulting intermonomer TM2-β3/β4 contact increases dimer stability. The entire TM2 helices are embedded in lipid acyl chains. Correspondingly, the cytoplasmic β-sheet is located near the start of the headgroup region in the cytoplasmic leaflet (Figures 8A and S11A). In contrast, the longer TM1 helices and the β4-TM2 linker push the periplasmic β-sheet toward the end of the headgroup region in the periplasmic leaflet. Whereas water is largely confined to only the lower face of the cytoplasmic β-sheet and solvates Lys23 and Lys25, it accesses both faces of the periplasmic β-sheet, in keeping with the fact polar sidechains project from both faces of this β-sheet. As expected, Arg19 and Arg92 snorkel into the headgroup region to hydrogen bond with lipid phosphates and water molecules.

Within each monomer, TM1 and TM2 pack tightly against each other. Between the two monomers, interhelical packing is generally tight but a fenestration occurs near the middle of the bilayer (Figure S11B). Interestingly, Gly44 and Ala78 are near this fenestration, so their mutation into a bulkier Val would improve the intermonomer helical packing and thereby increase dimer stability, consistent with our gel results (Figures 2 and S1). By contrast, Gly39 is away from any interhelical interface and its mutation to Val would not affect dimer stability, also in agreement with the gel results. Lastly, we added residues Met1-Ala16 to the dimer structure with these residues modeled as disordered and carried out MD simulations in the POPC:POPG membrane. In line with the PRE results (Figure 5), residues 1-10 exhibited high propensities of binding to the membrane (Figure S12).

## Discussion

Our considerable effort combining OS ssNMR, MAS ssNMR, and computational modeling and refinement has now produced a dimer structure for CrgA. This structure is very unusual, with a TM domain sandwiched between two β-sheets. The cytoplasmic and periplasmic β-sheets have much in common. They are both composed of one β-hairpin from each monomer; each β-hairpin has a rare type-VI turn with a Pro at the *i* + 2 position. The two β-sheets fully confine the TM2 helices but have missing corners at diagonal positions where the TM1 termini are located to provide linkages with the β-sheets. The short linkages on both sides of the membrane and interactions between the TM helices and the β-sheets keep the entire dimer structure rigid.

As demonstrated here, ssNMR is uniquely able to solve the structures of small membrane proteins and their complexes (< 30 kDa) in a membrane environment. These structures are too small for cryo-electron microscopy. Also, membranes are essential for their integrity and thus pose a formidable obstacle to crystallography ^27^. There are still only relatively few of these membrane protein structures in the PDB, making AlphaFold ^28^ predictions unreliable ^29^. The unusual structure of the CrgA dimer makes all these points especially pertinent.

With this dimer structure, we can begin to examine how CrgA interacts with its various partner proteins and regulates cell division (Figure 9A). The cytoplasmic β-sheet plus the TM1 N-termini (Figure 8B) represents a remarkable molecular platform in the membrane interfacial environment for protein interactions. This cytoplasmic surface of CrgA is positively charged, with a net charge of +6, and represents a favorable binding site for cytoplasmic folded domains or disordered regions with a negative charge. For example, although FtsQ can bind to CrgA via their TM helices, acidic residues are enriched in the disordered N-terminus (e.g., D_15_DAADEE_21_) ^30^ and thus favored to bind to the cytoplasmic surface of CrgA (Figure 9B). The folded GTPase domain of water-soluble FtsZ binds to an Arg/Ala-rich amphipathic α-helix of FtsQ ^30^, but FtsZ also has a disordered C-terminal tail enriched in acidic residues (e.g., D_367_DDDVD_372_), which may bind to the cytoplasmic surface of CrgA (Figure 9C). The binding with FtsZ localizes CrgA to the cell division site. It remains to be seen whether the periplasmic β-sheet of CrgA also serves as a binding site for other divisome proteins.

**Figure 9:**
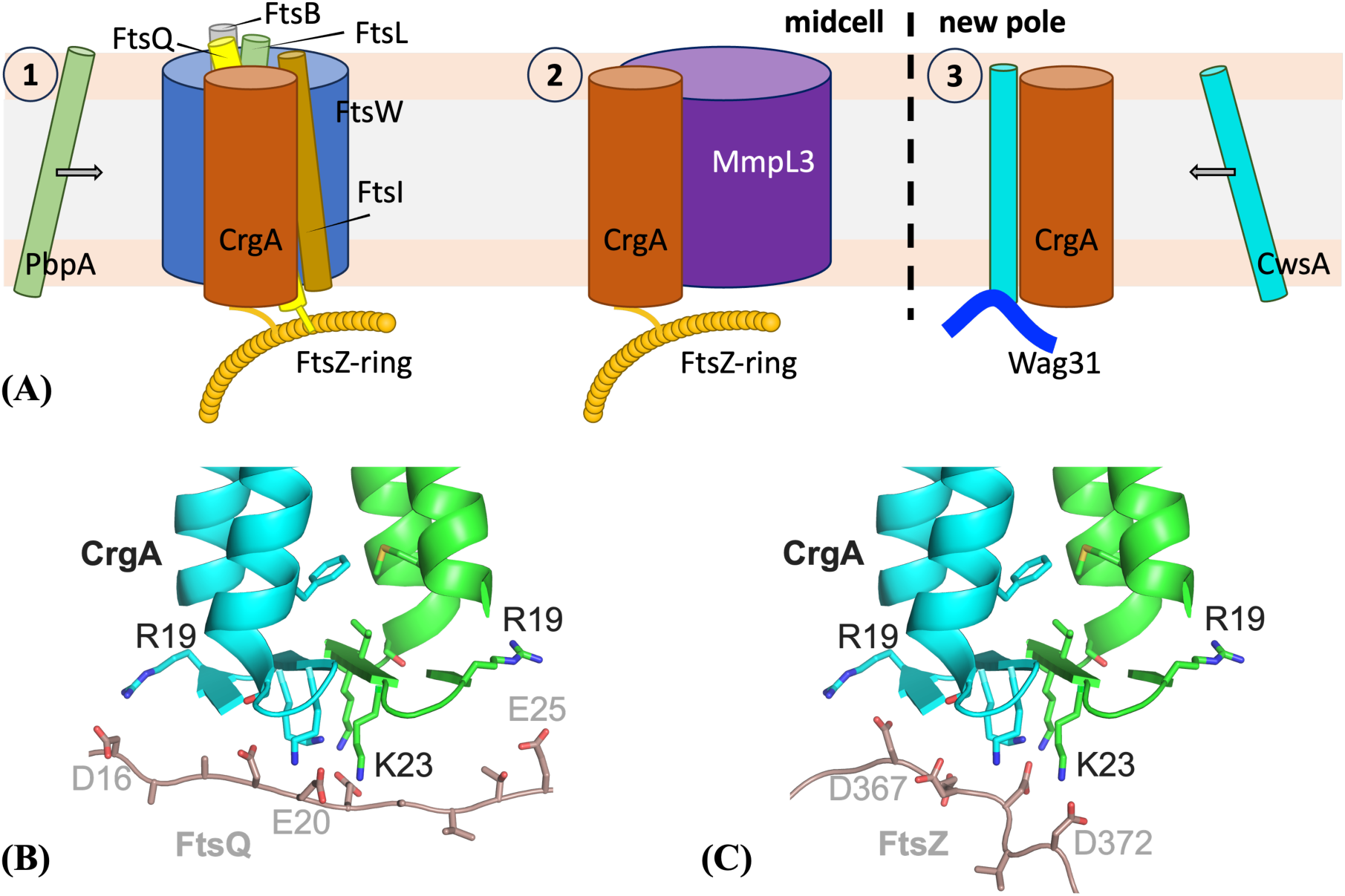
Interactions of CrgA dimer with other divisome proteins. (A) Roles of CrgA in three steps of *Mtb* cell division. (B) Binding of an acidic segment of FtsQ to the cytoplasmic β-sheet of CrgA. (C) Binding of an acidic segment of FtsZ to the cytoplasmic β-sheet of CrgA.

The dimerization of CrgA also generates a more substantial TM surface for interhelical binding with partner proteins. Of the five known TM binding partners ^16–18^, four (FtsQ, PBPA, FtsI, and CwsA) have only a single TM helix, whereas MmpL3 has 12 ^31^. For protein binding through TM domains, there is a distinct advantage if one of the proteins has more than a single helix, as partners can bind in its interhelical crevice or, more typically, bind across a pair of its helices. Both cases increase the contact surface and hence binding affinity. Moreover, for a protein such as CrgA that has multiple binding partners through this four-helix bundle, there are opportunities for more than one partner to be bound at the same time. The opportunities are further increased by the participation of the cytoplasmic β-sheet (or even the periplasmic β-sheet). For example, CrgA may bind with FtsZ via the cytoplasmic β-sheet and, at the same time, bind with FtsQ via the TM domain. Such simultaneous binding may help CrgA’s recruitment of the transpeptidases PbpA and FtsI to the nascent division site for peptidoglycan synthesis. Likewise, simultaneous binding of MmpL3 and other partners to CrgA can ensure coordinated construction of the different layers of the cell wall at the division site.

Lastly, the dimerization of CrgA allows it to bind two copies of CwsA at the same time. This binding facilitates the recruitment of Wag31 to the new cell pole for polar growth. Similar to FtsZ, Wag31 also self-assembles into filamentous structures ^32^. The binding of two copies of CwsA to CrgA increases the overall stability of the CrgA-CwsA-Wag31 ternary complex.

## Materials and Methods

### CrgA expression and purification

The crgA (Rv0011c) gene from *Mtb* H37Rv was cloned into a pET-29b vector (Novagen, Inc) including the C-terminal 6× His tag for protein purification. The cloned vector was transformed into *E. coli* DH5α cells (Stratagene, Inc). Amplified plasmids were purified with a QIAprep spin miniprep kit. Each pair of primers was designed and synthesized (Integrated DNA Technologies, Inc.) for PCR. In addition to the full-length CrgA protein, Δ1-29 truncated CrgA was prepared. For site-directed mutations, the whole plasmid with primers including the mutated site (e.g., G44V, G39V, N74A, or A78V) was used. To uniformly label with ^15^N or ^13^C, ^15^NH_4_Cl or ^13^C D-glucose was used as the sole nitrogen or carbon source in minimal media for protein expression. Specifically, for each 1 L of M9 medium, the following minimal salts were used: Na_2_HPO_4_ (6.8 g), KH_2_PO_4_ (3.0 g), NaCl (0.5 g), 1 M MgSO_4_ (2 mL), and 1 M CaCl_2_ (0.1 mL). This medium was then supplemented with 1 g ^15^NH4Cl and 2 g ^13^C glucose for uniform ^13^C and ^15^N labeling, respectively. For ^13^C- or ^15^N-amino acid specific labeled CrgA protein preparation, 20 amino acids (natural abundance) except the one intended to be ^15^N labeled were added to the 1 L M9 media. The amounts of the natural-abundance amino acids per L of M9 media were as follows: 800 mg of Asp and Glu, 500 mg of Ala, Val, Leu, Ile, and 200 mg of each of the remaining amino acids; the amount of the ^15^N-labeled amino acid was 200 mg. For ^15^N reverse labeled protein, the protocol was similar to that previously published ^33^. Protein purification followed protocols as reported ^16^ and later in more detail ^19^. Cells were thawed in a 37 °C water bath. 1 μl of benzonase nuclease was added to cells and incubated in ice for 30 minutes before cell lysis. Cells were lysed three times using a French press (Thermo Scientific, Inc) at 10,000 PSI to disrupt the cell wall and membrane. The lysate was pelleted down after centrifuging (8,000 g for 20 minutes at 8 °C), and the inclusion body was separated from the supernatant and resuspended with solubilization buffer (50 mM Tris buffer pH 8.0 with 200 mM NaCl and 3% Empigen). The supernatant was separately ultra-centrifuged at 228,000 g for 1.5 hours to obtain the membrane fraction. The membrane fraction was bath sonicated for 10-15 minutes at room temperature and solubilization buffer was added. Both solutions containing inclusion bodies and membrane fraction were incubated to solubilize proteins in an orbital shaker at 4 °C overnight. Prior to protein purification, the mixture was centrifuged at 228,000 g for 30 minutes to remove any insoluble materials. Before loading the supernatant, a 5 ml His-Prep fast flow Ni^2+^ affinity column (GE Lifesciences, Inc) was equilibrated with solubilization buffer in an AKTA Xpress system (GE Lifesciences, Inc). Imidazole was added to the supernatant to reach a final concentration of 20 mM, and the mixture was loaded onto the Ni^2+^ affinity column. The column was washed with wash buffer 1 (50 mM Tris buffer pH 8.0 with 200 mM NaCl, 2% Empigen, and 20 mM imidazole) to remove most of the impurities in the column until the UV absorbance reached baseline. High imidazole concentration wash buffer 2 (50 mM Tris buffer pH 8.0 with 200 mM NaCl, 0.7% Empigen, and 60 mM imidazole) was flowed into the column to achieve higher purity (with some loss in CrgA). The column was washed with wash buffer 3 (50 mM Tris buffer pH 8.0 with 200 mM NaCl and 0.4% DPC) until the UV absorbance reached baseline. CrgA was finally eluted with elution buffer (50 mM Tris buffer pH 8.0 with 200 mM NaCl, 0.4% DPC, and 200 mM imidazole). Typically, the yield of CrgA was 20-30 mg from 1 L of M9 media.

### CrgA reconstitution into POPC:POPG lipid bilayers

POPC/POPG (4:1 molar ratio) liposomes were chosen to mimic the highly negatively charged inner membranes of mycobacteria ^23^. POPC and POPG were purchased from Avanti Polar Lipids. A lipid film was obtained by evaporating chloroform from stock lipid solutions using N_2_ gas, followed by vacuum drying overnight. The film was solubilized with 5 mM Tris buffer (pH 8.0), including 0.2% DPC, and the lipid solution was sonicated for 10 minutes. Methyl-β-cyclodextrin was used to remove DPC and facilitate the incorporation of CrgA into liposomes ^34^. The CrgA monomer-to-lipid ratios were 1:80 for all OS experiments and 1:40 for MAS experiments. For PRE experiments, proteoliposome samples were doped with 0.5%, 1%, and 2% (mol/mol) of 16:0 PE-DTPA (Gd^3+^).

### OS ssNMR

All OS ssNMR experiments were performed on a Bruker Avance 600 MHz NMR spectrometer. The protocol for uniform alignment of the lipid bilayer preparations of membrane proteins has been described in detail ^34^. Briefly, lipid bilayer preparations are spread on thin glass slides and then the glass slides are stacked in a short segment of square glass tubing inserted into a solenoid coil perpendicular to the magnetic field of the NMR. OS spectra were acquired using the PISEMA pulse sequence ^35^ at 13 °C with a home-built low-E static NMR probe ^36^. ^15^N chemical shifts were referenced to the ^15^N signal of an aqueous solution of ^15^N-labeled ammonium sulfate (5%, pH 3.1) at 26.8 ppm. The typical acquisition protocol involved ^1^H 90° pulse length of 4 μs, ^1^H and ^15^N RF fields of 50 kHz, ^1^H decoupling RF field of 62.5 kHz, recycle delay of 4 s, cross-polarization contact time of 1000 μs with 2000 scans, and 28-32 increments in the dipolar coupling dimension with 4000-5000 scans for each increment. The orientational restraints were interpreted using a motionally averaged chemical shift tensor (σ_11_ = 57.3, σ_22_ = 81.2, and σ_33_ = 227.8 ppm) and a motionally averaged ^15^N-^1^H dipolar interaction magnitude of 10.735 kHz. The relative orientation of σ_33_ and the ν_||_ component of the dipolar interaction was set at 17° ^37^.

### MAS ssNMR

For MAS experiments, proteoliposome samples were evenly packed in the MAS rotor using a swinging bucket rotor. All ^13^C chemical shifts were referenced externally to the ^13^C carboxyl resonance of glycine at 178.4 ppm. 1D CP spectra were acquired using a Bruker Avance 600 MHz spectrometer with a home-built, low-E ^1^H-^13^C double resonance probe ^38^. RF field strengths were 75-80 kHz for ^1^H and 70-75 kHz for ^13^C. 1 ms of contact time and a ^1^H RF field of 66 kHz were set for the ^1^H-^13^C CP condition. 75-80 kHz ^1^H SPINAL64 decoupling was applied during acquisition. The same parameters from the 1D experiment were used for 2D ^13^C-^13^C correlation experiments through dipolar couplings. Mixing times were varied (50 ms to 800 ms) depending on the distance between different ^13^C sites. 1D INEPT spectra were acquired using a Bruker Avance III HD 800 MHz spectrometer with a home-built ^1^H-^13^C-^15^N triple resonance probe. The rotor was spun at 12 kHz. RF field strengths were 70-72 kHz for both ^1^H and ^13^C. The rotor periods of ^1^H and ^13^C evolution were optimized to maximize signals. 70-72 kHz ^1^H SPINAL64 decoupling was applied during acquisition. The same parameters from the 1D experiment were used for 2D ^13^C-^13^C correlation experiments through J-couplings. The number of rotor periods was optimized to enhance TOBSY mixing.

### Modeling of monomeric TM domain and dimeric TM domain

We represented TM1 (residues Ser29-Ala55) and TM2 (Trp73-Trp92) as Cα-only ideal helices to explore the full range of models that satisfied the OS and DARR restraints. The ideal helices had standard geometries, including a radius of 2.3 Å, a rise of 1.5 Å per residue, and a rotation of 100° per residue. The tilt angles of both helices were fixed to 13°, as determined previously ^19^.

For modeling the monomeric TM domain, we defined the center of the cross-section at Met41 of TM1 as the center of this helix; similarly, the center of TM2 was defined using Ala80. Initially, both helices were aligned parallel to the *z*-axis; TM1 had its center fixed at the origin, whereas the center TM2 was restricted to an elevation of 3 Å (corresponding to a rise of two helical residues). After choosing the self-rotation of TM1 to have Met41 on the negative *x*-axis, TM1 was tilted around the *x*-axis (for 13°) and then fixed in space. TM2 had four degrees of freedom: the *x* and *y* coordinates of its center, the self-rotation, and the azimuthal angle of its helical axis. We scanned the Cartesian coordinates at 2-Å intervals and the angles at 30° intervals. Monomer models that satisfied the two intramonomer DARR restraints, between Met41 and Ala80 and between Leu48 and Tyr75, and avoided TM1-TM2 clashes were selected. For the Met41-Ala80, the upper bound for Cα-Cα distance was 8 Å; for the Leu48-Tyr75, this bound was increased to 10 Å, to account for the fact that one of the DARR partner sites was a sidechain atom (i.e., Leu48 Cγ) instead of Cα. Clashes were detected using a 5.1 Å cutoff for any TM1-TM2 Cα-Cα distance ^39,40^. Each selected model was replaced by an all-atom model, again representing TM1 and TM2 as ideal helices, and aCS and DC were back-calculated to find acceptable models for the monomeric TM domain, which satisfied all DARR and OS restraints.

The acceptable models for the monomeric TM domain were used to generate models for the dimeric TM domain. There were two additional degrees of freedom, namely the *x* and *y* coordinates of the C2 symmetry axis. These coordinates were again scanned at 2-Å intervals.

Those satisfying the intermonomer Phe33-Met90 DARR restraint (Cα-Cα distance < 10 Å) and avoiding intermonomer clashes were identified as acceptable models for the dimeric TM domain. Finally, a single model survived based on filtering using Ser29-Ser29 and Ala55-Ala55 Cα-Cα distances; the surviving model had these distances at 18.8 and 17.5 Å, respectively. This surviving model was improved slightly by a finer scan, with Cartesian coordinates at 1-Å intervals and the angles at 10° intervals. The Cα model was replaced by an all-atom model for the next step, with sidechains adjusted manually to better satisfy DARR restraints for the Met41-Ala80 and Leu48-Tyr75 pairs.

### Generation of dimer model for residues Val17-His93

The cytoplasmic and periplasmic β-sheets were built using Xplor-NIH ^41^ by simulated annealing with C2 symmetry imposed. The cytoplasmic β-sheet consisted of Val17-Gly27, with Val17-Thr20 and Met22-Gly27 as β-strand residues. The type-VI turn centered at Thr20-Pro21 was modeled after PDB entry 1TMN residues T_49_LPG_52_. Intramonomer hydrogen bonds were formed between Val17 CO and Val24 NH and between Arg19 and Met22; intermonomer hydrogen bonds were formed between Pro21 CO and Gly27 NH and between Lys23 and Lys25 (Figure S8). Similarly, the periplasmic β-sheet consisted of Ala55-Met67, with Ala55-Gln59 and Thr62-Met67 as β-strand residues and Ala60-Pro61 forming a type-VI turn. Intramonomer hydrogen bonds were formed between Ala55 CO and Trp66 NH, between Gly57 and Leu64, and between Gln59 and Thr62; intermonomer hydrogen bonds were formed between Ala63 and Met67 and between Asn65 and Asn65 (Figure S9). The (ϕ,ψ) angles of β-strand residues were restrained to (140°, 130°); hydrogen bonding atoms were also distance-restrained (3 Å for O-N and 2 Å for O-H). Simulated annealing was performed with the temperature ramping down from 3500 K to 25 K in 1000 steps, with the force constants ramping up from 5 to 1000 kcal/mol rad^-2^ for angle restraints and from 2 to 30 kcal/mol Å^-2^ for distance restraints.

Using VMD ^42^, the surviving model for the dimeric TM domain (with His93 appended to TM2) was aligned with the cytoplasmic and periplasmic β-sheets as illustrated in Figures S8 and S9, respectively. The TM domain and the two β-sheets were joined using MODELLER (version 9.22 ^43^) to build the initial dimer structure for residues Val17-His93.

### Refinement of the dimer structure

Refinement was carried out by running restrained MD simulations in NAMD ^44^ with the CHARMM36 force field for protein and lipids ^45^ and TIP3P for water ^46^. The initial dimer structure was placed in a POPC:POPG (4:1) bilayer (125 lipids per leaflet) and solvated with 16589 water molecules and Na^+^ and Cl^-^ ions for charge neutralization and a 150 mM salt concentration in a box with dimensions of 95.5 × 95.5 × 101.4 Å^3^. The total number of atoms was 85531. After 10000 cycles of conjugate-gradient energy minimization, six steps of equilibration were performed, with constant NVT for the first two steps and constant NPT for the last four steps and with durations of 125, 125, 125, 500, 500, 500 ps, respectively. The timesteps were 1 fs in the first three steps and 2 fs in the last three steps. Harmonic restraints on lipid headgroups were gradually reduced from 5 to 0 kcal mol^-1^ Å^-2^ and positional restraints on protein backbone heavy atoms were reduced from 10 to 1 kcal mol^-1^ Å^-2^. The (ϕ,ψ) angles of β-strand and α-helical residues were restrained to (140°, 130°) and (−60°, −45°), respectively, with a force constant of 100 kcal mol^-1^ rad^-2^. Hydrogen bond distances in β-sheets and α-helices were restrained with force constants of 50 and 20 kcal mol^-1^Å^-2^, respectively.

After equilibration, OS restraints were gradually introduced by increasing the force constants to 3.0 kcal mol^-1^ kHz^2^ for DC and 0.02 kcal mol^-1^ ppm^2^ for aCS over 500 ps at constant NPT. Additional restraints included C2 symmetry (force constant at 5 kcal mol^-1^ Å^-2^ between actual and symmetry-predicted Cα positions) and DARR restraints (total of 36 distances, each restrained to be within 8 Å with a force constant of 100 kcal mol^-1^Å^-2^. The simulation was continued for another 500 ps. In the third 500-ps simulation, (ϕ,ψ) angle restraints were removed for residues 51-55. This 500-ps segment was repeated to generate 10 models for deposition to the PDB. The timestep was 1 fs in these restrained simulations. OS restraints were implemented using a TCL force interface with NAMD.

In all the simulations, the particle mesh Ewald method ^47^ was used to treat long-range electrostatic interactions. The cutoff distance for nonbonded interaction was 12 Å, with van der Waals interactions force switching at 10 Å. All bonds involving hydrogen atoms were constrained by the SHAKE algorithm ^48^. Temperature was maintained at 310 K using the Langevin thermostat with a damping constant of 1 ps^-1^; pressure was maintained at 1 atm by the Langevin piston method ^49^.

### MD simulations of full-length CrgA dimer in membranes

A full-length CrgA dimer was built from model 1 (residues V17-H93) of the refinement simulations. Residues Met1-Ala16 were modeled as disordered. The full-length CrgA dimer was placed in a POPC:POPG (4:1) bilayer (200 lipids per leaflet), solvated with 36303 water molecules and 150 mM NaCl, resulting in a total of 165227 atoms in a box with dimensions of 119.0 × 119.0 × 124.9 Å^3^. The energy minimization and equilibration were performed as for the preceding refinement simulations. In the next 2 ns of simulation at constant NPT, the (ϕ,ψ) angles of β-strand and α-helical residues were restrained with force constants at 100 and 50 kcal mol^-1^ rad^-2^, respectively. In the final production run of 100 ns at constant NPT, the (ϕ,ψ) restraints on β-strand residues were reduced to 50 kcal mol^-1^ rad^-2^. The timesteps were 2 fs for the last two runs. Snapshots were saved at 100 ps intervals for analysis. Membrane contact probabilities of individual residues were calculated from the last 40 ns of the simulation. For each snapshot, a residue was defined as membrane-contacting if any of its heavy atoms came within 3.5 Å of a lipid heavy atom. The results were averaged over 400 snapshots and then over the two monomers.

## Acknowledgments

This work was supported by National Institutes of Health Grants AI119178, GM122698, GM148766, and GM118091. All NMR experiments were carried out at the National High Magnetic Field Laboratory supported by the NSF Cooperative Agreements DMR1644779 and DMR2128556 and by the State of Florida.

## Author contributions

T.A.C. and H.-X.Z. designed the research; Y.S., R.P., N.D., J.T., H.Q., M.H., Y.-Y.H., R.F., R.Z., H.-X.Z., and T.A.C. performed the research and analyzed the data; H.-X.Z., T.A.C., R.Z., and R.P. wrote the manuscript.

## Notes

### Competing Interest Statement

The authors have declared no competing interest.

### Summary of Updates

The acknowledgments section is updated to include one more grant.

