## Supporting Figures for "*Mycobacterium tuberculosis* CrgA Forms a Dimeric Structure with Its Transmembrane Domain Sandwiched between Cytoplasmic and Periplasmic β-Sheets, Enabling Multiple Interactions with Other Divisome Proteins"

*Supporting Information*

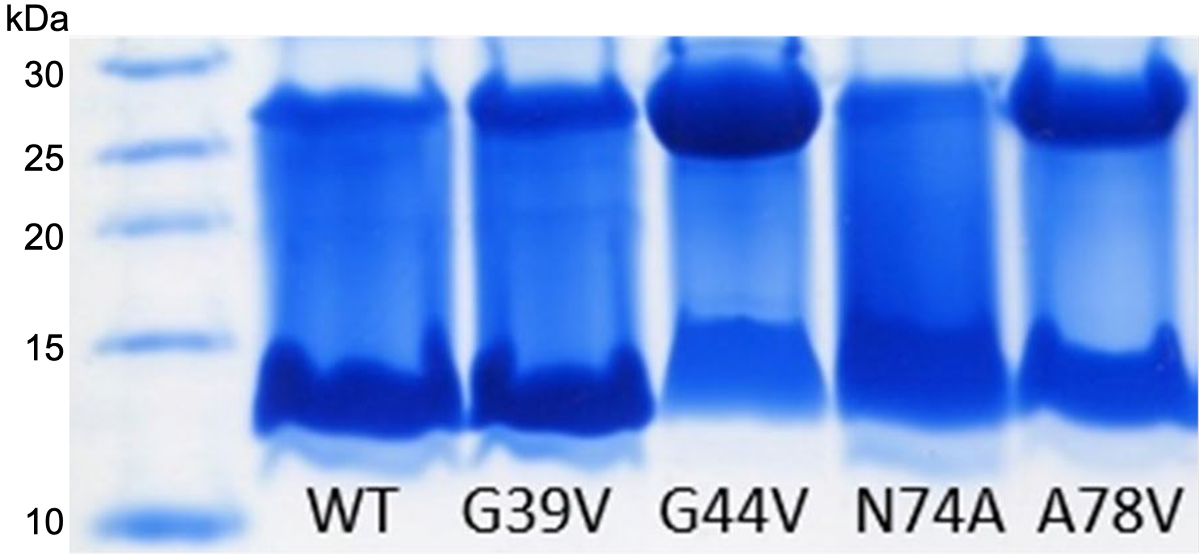


Figure S1. A 12% SDS-PAGE gel of CrgA variants. Compared to the wild-type (WT) protein, a more intense dimer band is only shown by the G44V and A78V mutants but not the G39V and N74V mutants. The molecular weights of monomer and dimer are 11.5 kDa and 23 kDa, respectively.

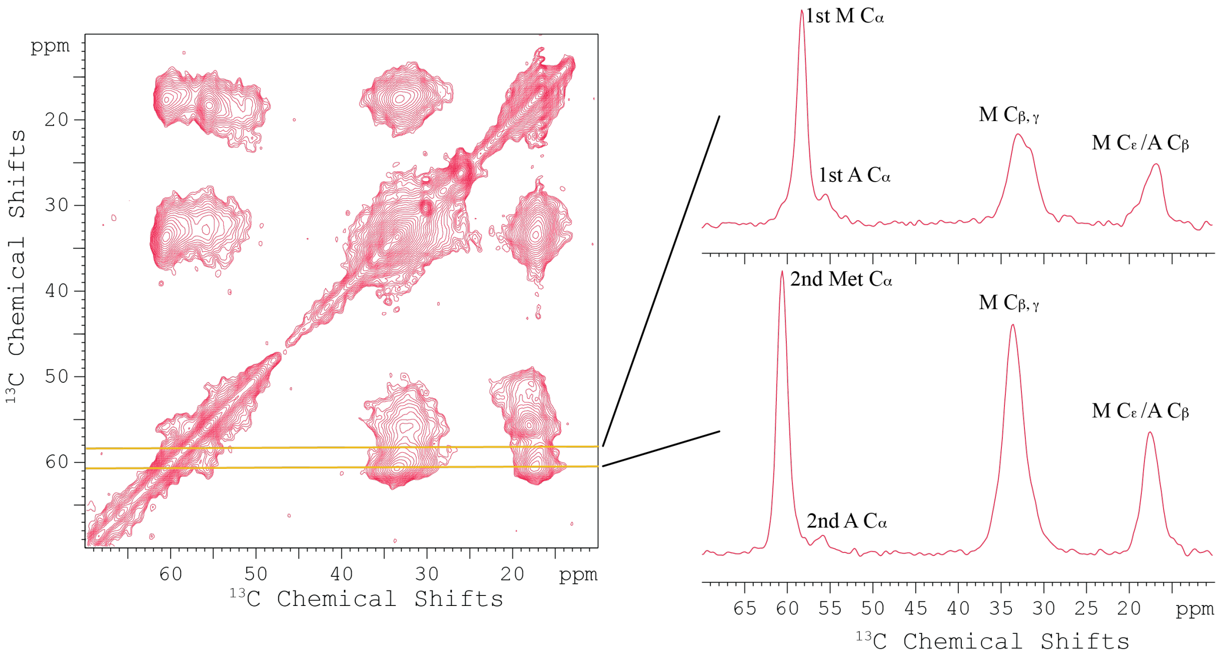


Figure S2. 2D DARR $^{13}C$-$^{13}C$ correlation spectrum of $^{13}C$-Ala/Met labeled CrgA in POPC:POPG membranes at 265 K. Spectral slices through two Met Cα frequencies, at 58.3 ppm and 60.8 ppm, are displayed on the right. Collected at 600 MHz with a 12.2-kHz spinning rate.

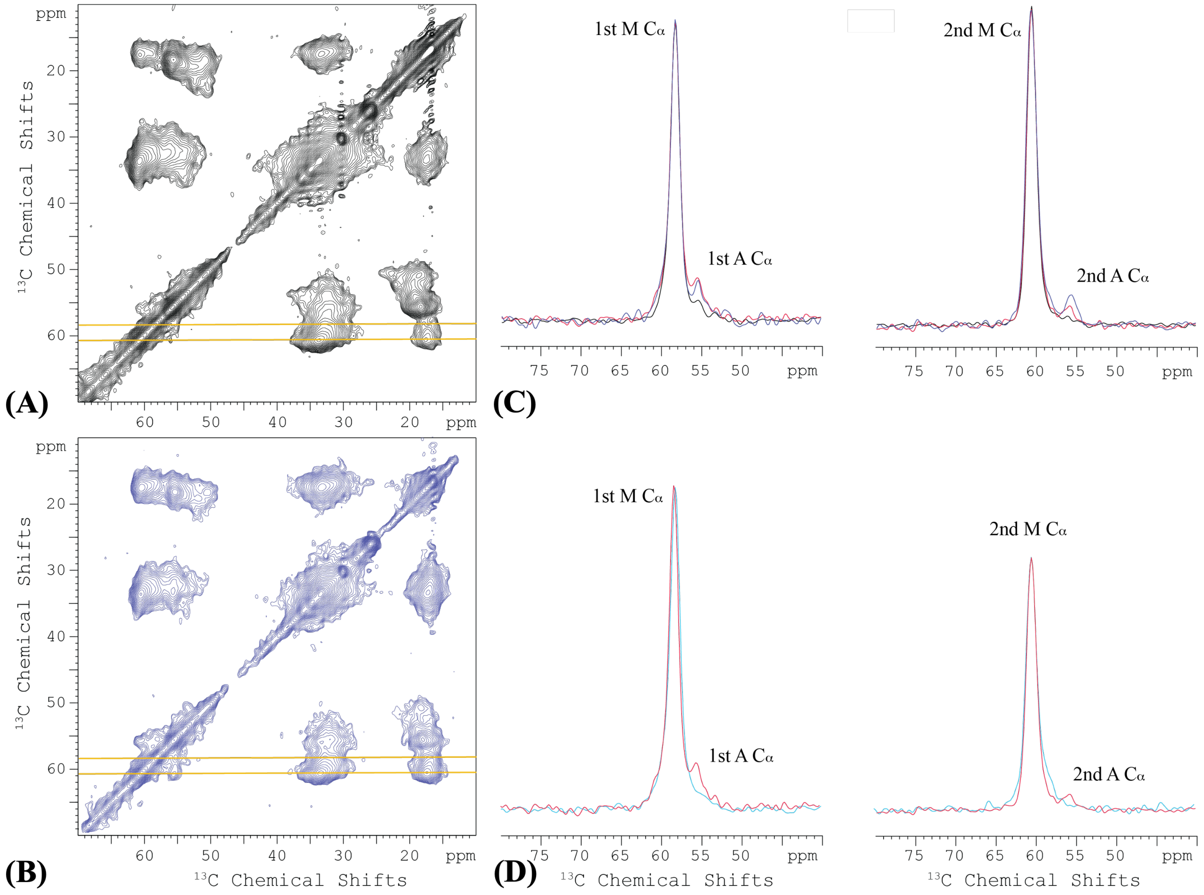


Figure S3. 2D DARR $^{13}$C-$^{13}$C correlation spectra of $^{13}$C-Ala/Met labeled WT CrgA and the A78V/A80G double mutant in POPC:POPG membranes at 265 K. (A) WT at a 100 ms mixing time. (B) WT at a 800 ms mixing time. (C) Spectral slices through Met Cα frequencies at 58.3 ppm (left) and 60.8 ppm (right) from 100 (black), 500 (red), and 800 (blue) ms mixing times. (D) Overlay of spectral slices from 2D $^{13}$C-$^{13}$C correlation spectra of WT (red) and A78V/A80G (light blue) at a 500 ms mixing time. In panels (C) and (D), the diagonal peak intensities were scaled to the same height for better comparison of off-diagonal peaks.

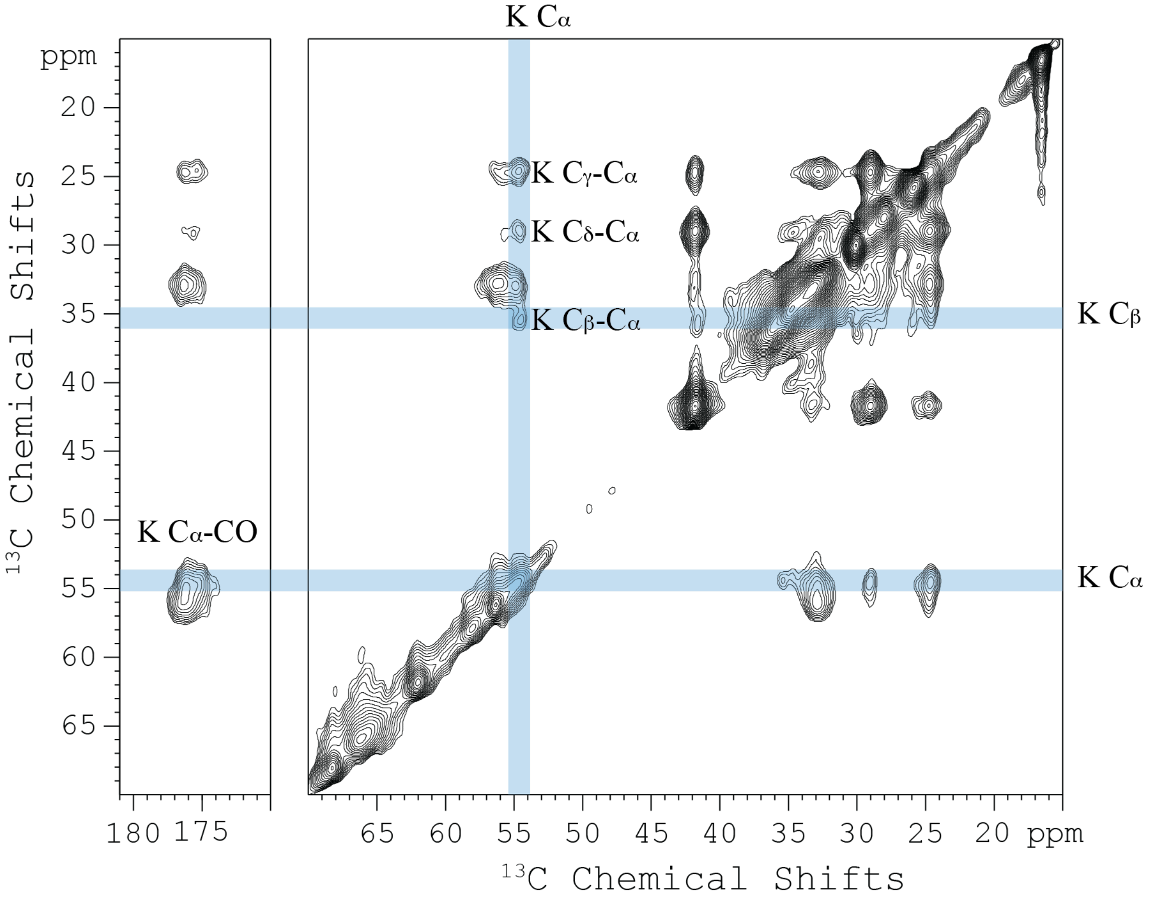


Figure S4. 2D DARR $^{13}C$-$^{13}C$ correlation spectrum of $^{13}C$-Lys labeled CrgA in POPC:POPG membranes at 265 K. Collected at 600 MHz with a 100 ms mixing time and 12.2-kHz spinning rate.

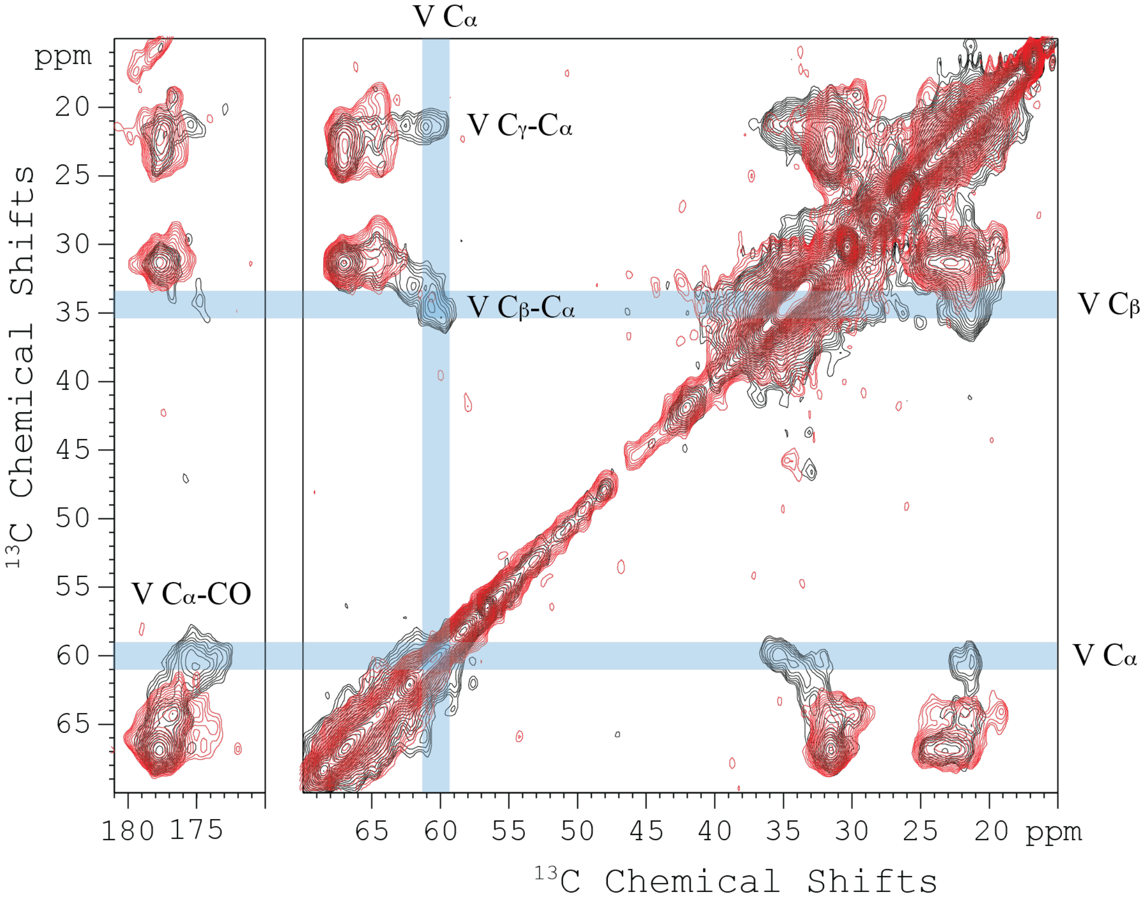


Figure S5. 2D DARR $^{13}C$-$^{13}C$ correlation spectra of $^{13}C$-Val labeled WT (black) and Δ2-28 (red) CrgA in POPC:POPG membranes at 265 K. Collected at 600 MHz with a 100 ms mixing time and 12.2-kHz spinning rate.

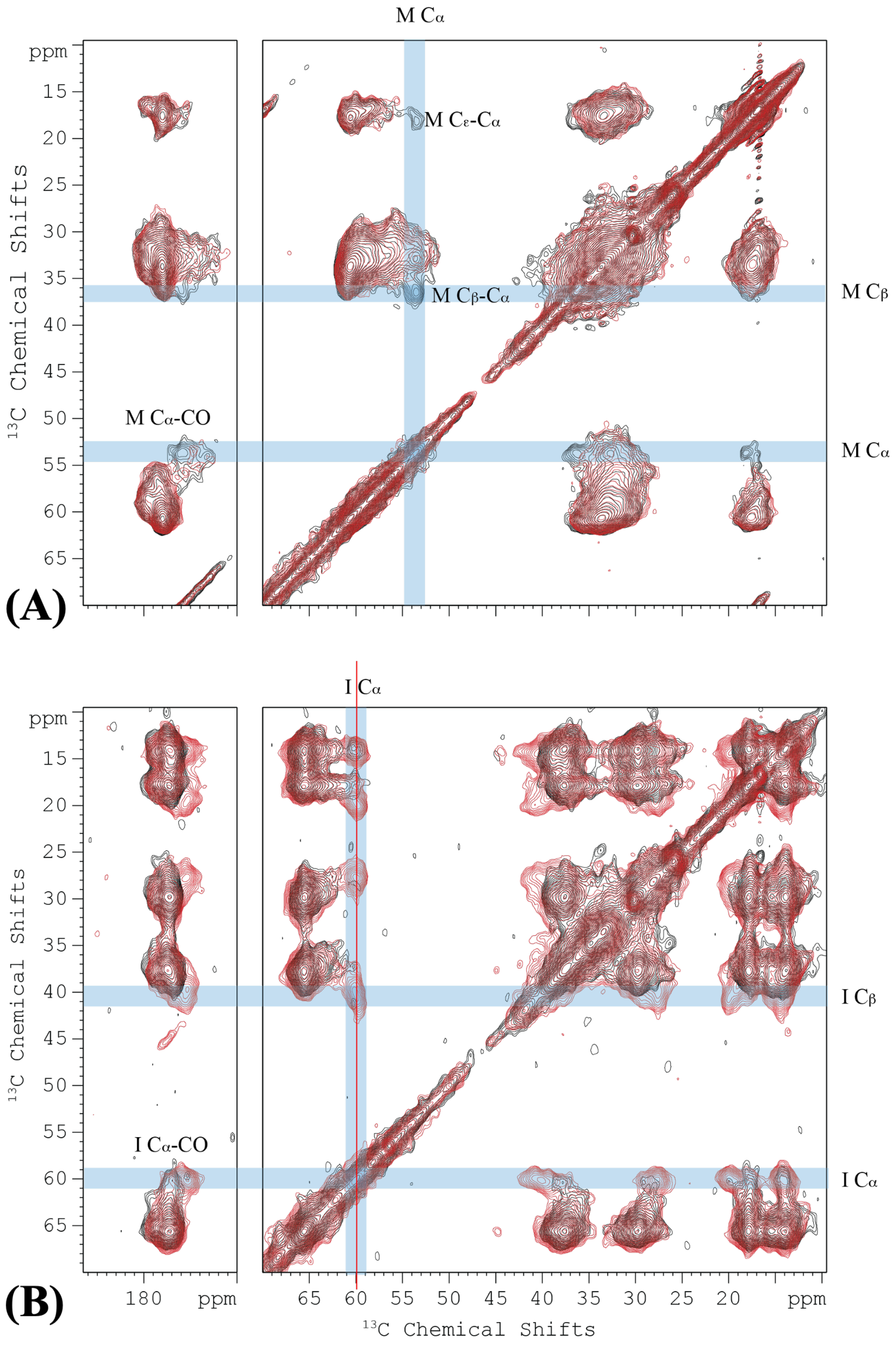


Figure S6. 2D DARR $^{13}$C-$^{13}$C correlation spectra of four $^{13}$C-labeled samples in POPC:POPG membranes at 265 K. (A) $^{13}$C-Met labeled WT (black) and T20A/M22I (red), with an 8-kHz spinning rate. (B) $^{13}$C-Ile labeled WT (black) and T20A/M22I (red), with a 10-kHz spinning rate. Collected at 600 MHz with a 100 ms mixing time.

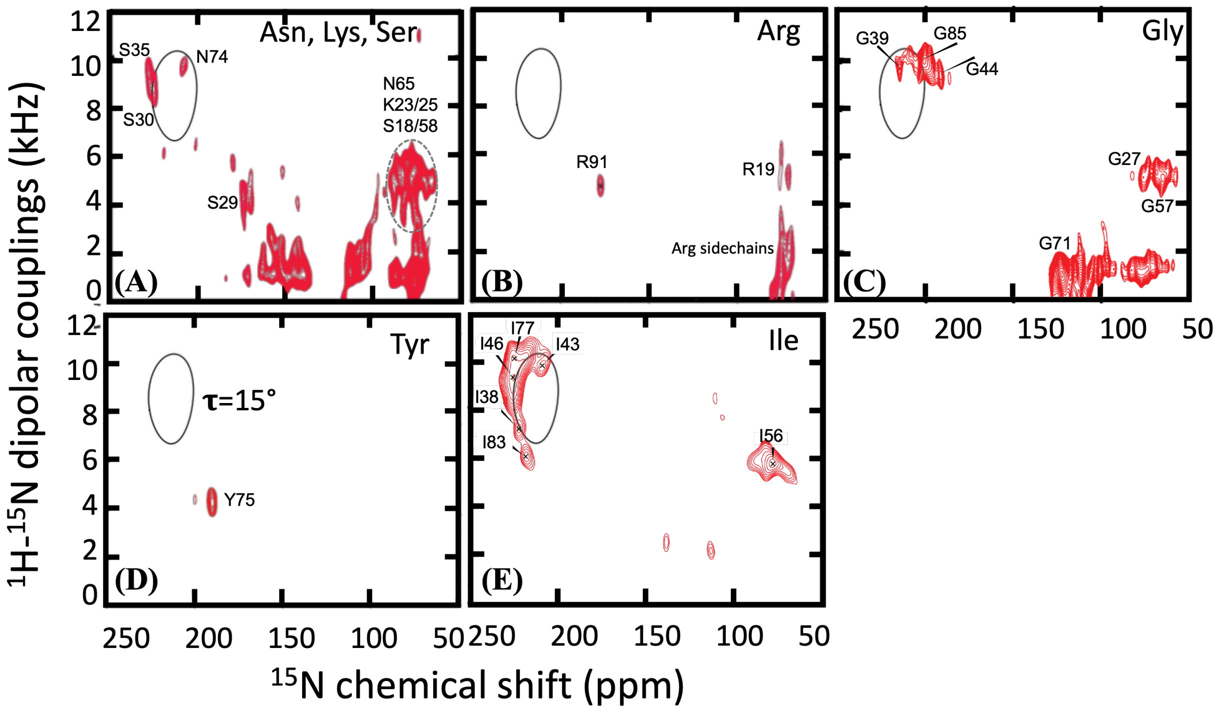


Figure S7. OS ssNMR spectra of amino-acid-specific $^{15}$N-labeled samples. A PISA wheel for a TM helix with a 15° tilt angle is shown in solid curve. (A) Reverse $^{15}$N-labeled Asn/Lys/Ser. (B) Reverse $^{15}$N-labeled Arg. (C) $^{15}$N-Gly. (D) $^{15}$N-Tyr. (E) $^{15}$N-Ile. The helical portion of Panel (E) was published previously in ref 19.

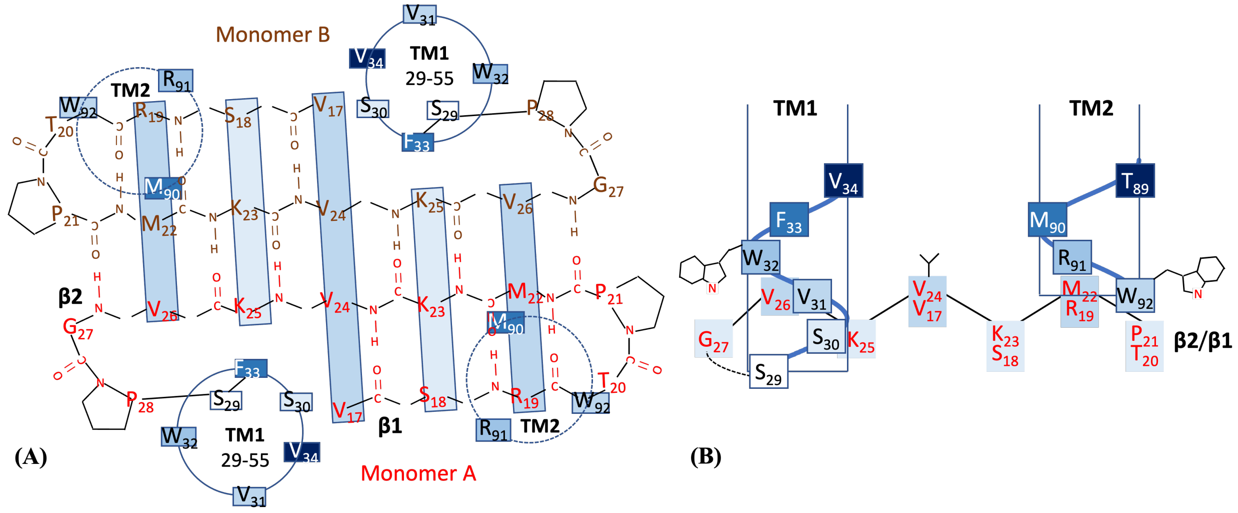


Figure S8. Illustration of the cytoplasmic β-sheet and the positioning of the TM helical termini relative to it. (A). Bottom view. (B) Side view. Blue colors in light to dark shades indicate increasing elevation toward the bilayer middle. In (A), the two faces of the β-sheet are indicated by boxes that are drawn through Cα atoms and shaded in lighter (water side) and darker (membrane side) blue.

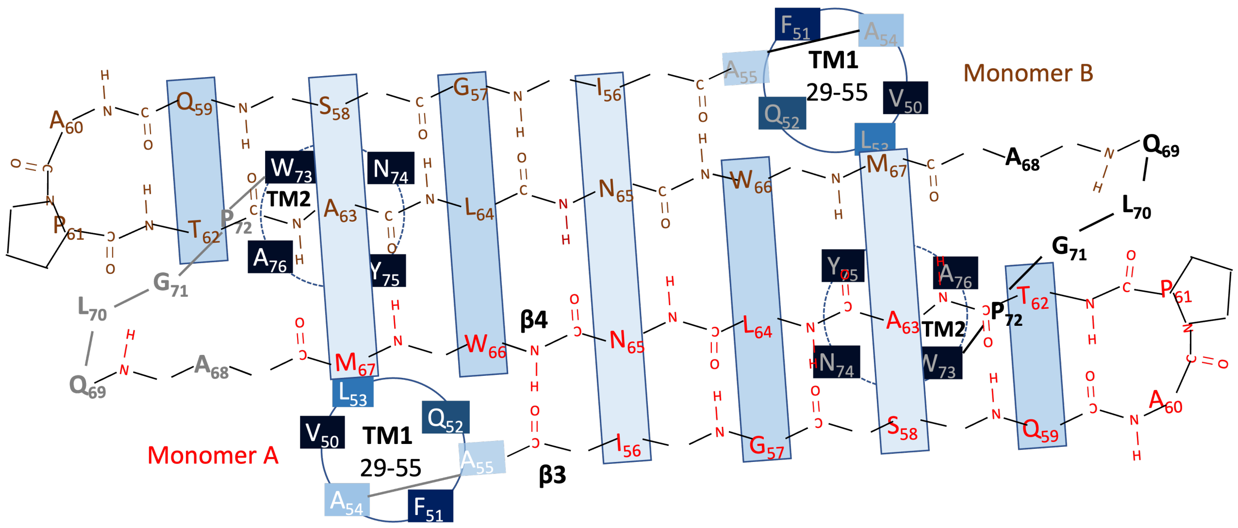


Figure S9. Illustration of the periplasmic β-sheet and the positioning of the TM helical termini relative to it. Blue colors in light to dark shades indicate increasing recession toward the bilayer middle. The two faces of the β-sheet are indicated by boxes that are drawn through Cα atoms and shaded in lighter (water side) and darker (membrane side) blue.

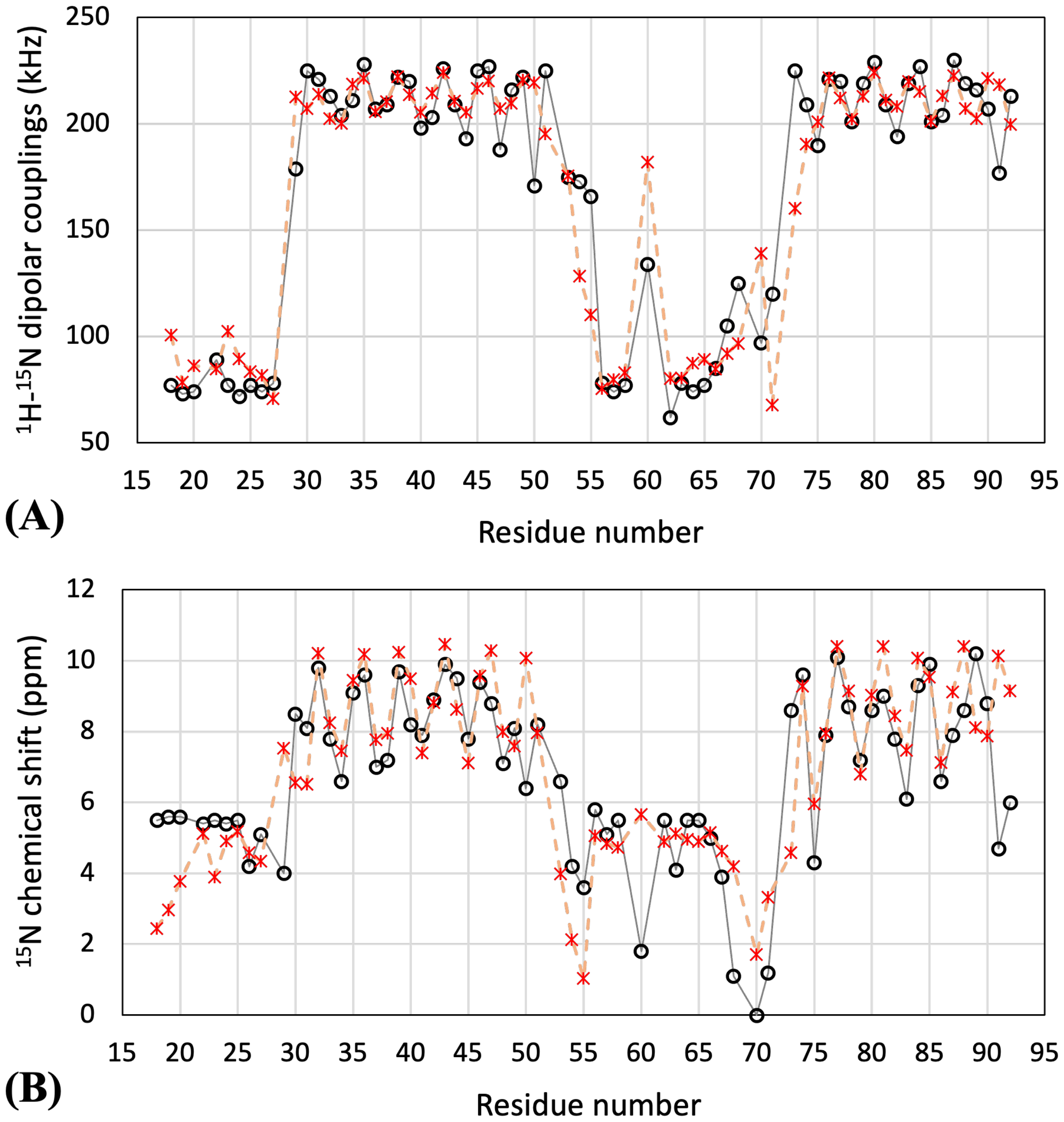


Figure S10. Comparison of observed (circles) and back-calculated (asterisks) OS properties. (A) Dipolar couplings. (B) Anisotropic chemical shifts.

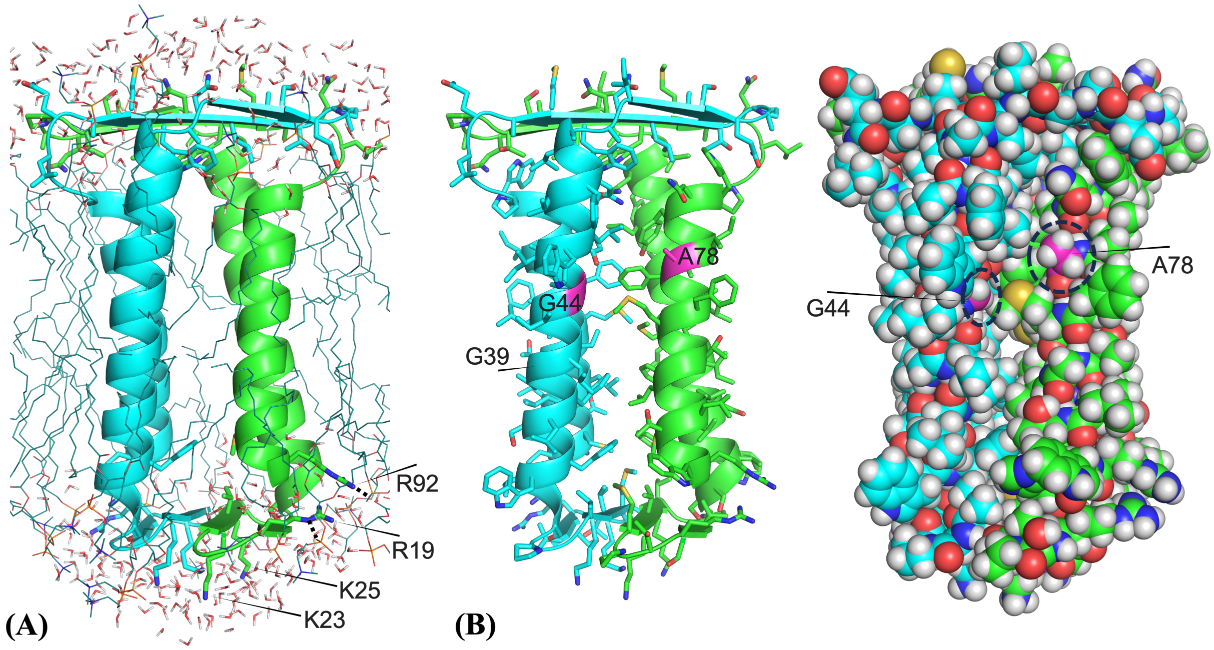


Figure S11. Membrane and water interactions and packing of CrgA dimer. (A) Membrane and water interactions. Arg19 and Arg92 form salt bridges (indicated by dashed lines) with lipid phosphates, whereas Lys23 and Lys25 are solvated by water. (B) Packing between the TM helices of the two monomers, shown by a stick representation of sidechains (left) and a sphere representation of all atoms (right). A fenestration occurs near the middle of the TM domain. Gly44 and Ala78 line this fenestration, whereas Gly39 faces lipid acyl chains. The view in this figure is rotated around the vertical axis by 30° from that in Figure 8A.

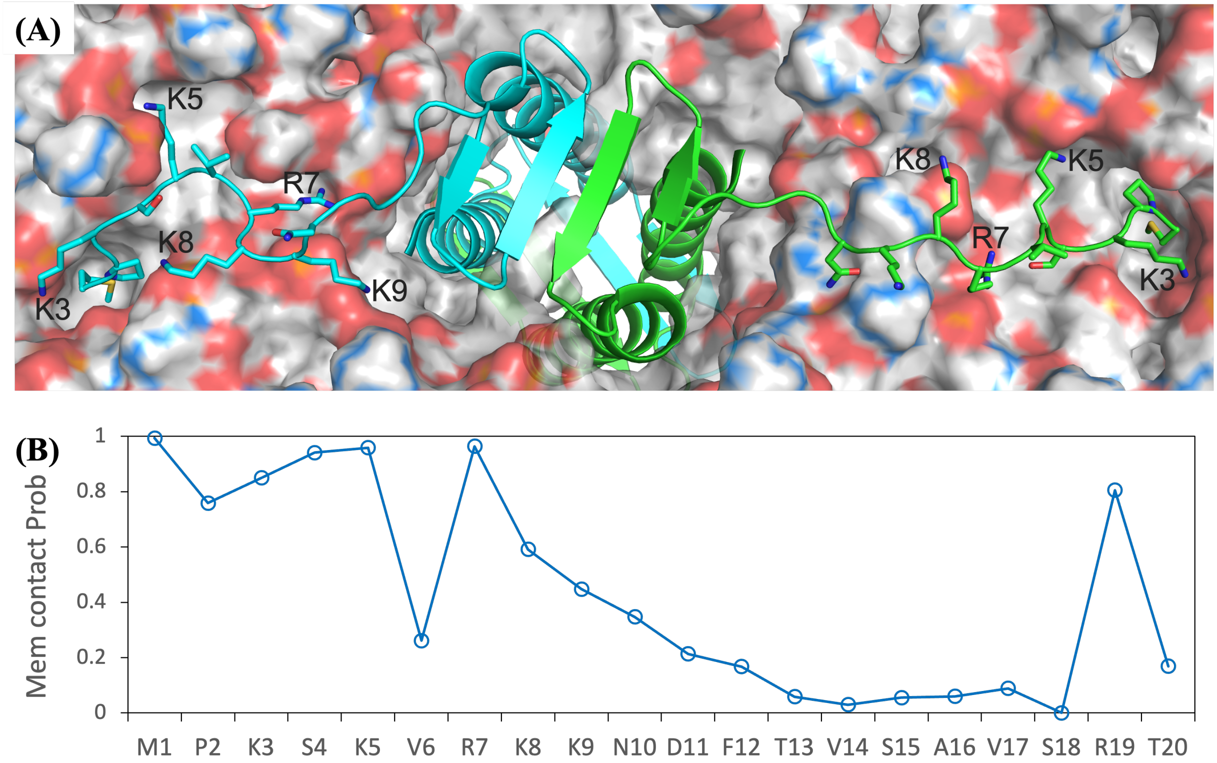


Figure S12. The N-terminal disordered region shows significant propensities for membrane association in MD simulations of full-length CrgA dimer in a POPC:POPG membrane. (A) A snapshot from the simulations. The membrane is displayed as surface, with oxygen, phosphorus, nitrogen (not visible), hydrogen, carbon on cholines, and other carbon atoms in red, orange, blue, white, light blue, and gray, respectively. (B) Membrane contact probabilities of the first 20 residues.
